# Polyglutamylation of microtubules drives neuronal remodeling

**DOI:** 10.1101/2024.03.11.584412

**Authors:** Antoneta Gavoci, Anxhela Zhiti, Michaela Rusková, Maria M. Magiera, Mengzhe Wang, Karin A. Ziegler, Torben Johann Hausrat, Stefan Engelhardt, Matthias Kneussel, Martin Balastik, Carsten Janke, Thomas Misgeld, Monika S. Brill

## Abstract

Developmental remodeling shapes neural circuits via activity-dependent pruning of synapses and axons. The cytoskeleton is critical for this process, as microtubule loss via enzymatic severing is an early step of pruning across many circuits and species. However, how microtubule-severing enzymes, such as spastin, are activated in specific neuronal compartments remains unknown. Here, we reveal that polyglutamylation, a posttranslational tubulin modification that is enriched in neurons, plays an instructive role in developmental remodeling by tagging microtubules for severing. Motor neuron-specific gene deletion of enzymes that add or remove tubulin polyglutamylation—TTLL glutamylases vs. CCP deglutamylases—accelerates or delays neuromuscular synapse remodeling in a neurotransmission-dependent manner. This mechanism is not specific to peripheral synapses but also operates in central circuits, e.g., the hippocampus. Thus, tubulin polyglutamylation acts as an activity-dependent rheostat of remodeling and shapes neuronal morphology and connectivity.

## Introduction

Microtubules are regulators of cellular shape, dynamics, and transport—processes that are especially difficult to coordinate in the complex cells of the nervous system ^1–3^. As a consequence, microtubule functions in neurons need to be locally regulated. In analogy to how posttranslational modifications (PTMs) diversify histone function, the concept of the “tubulin code” posits that microtubules can be endowed with unique functions through PTMs and a set of proteins that add, remove, or interpret them. Such a system of “writer”, “eraser”, and “reader” microtubule-associated proteins (MAPs) and enzymes can then locally modify microtubule stability, as well as the trafficking characteristics and cargo selection of intracellular transport with high spatiotemporal precision ^3–5^. Thus, in principle, the tubulin code can explain how a general system of filamentous proteins could locally regulate a plethora of parallel functions to meet the compartmentalized needs of neural cells. Indeed, the microtubular cytoskeleton of neurons is very specialized. This specialization includes the expression of neuron-specific tubulin isoforms (such as tubulin beta-3, Tubb3), pronounced and compartmentalized PTMs, as well as a unique set of MAPs and severing enzymes that play important roles in neurodegenerative and neurodevelopmental diseases ^6–10^. Nevertheless, while a PTM-based implementation of the “tubulin code” is an appealing concept and clearly plays important roles in the basic health and cell biology of neurons, e.g. via cell migration, axonal transport, growth cone navigation, and cilia function ^11^, how the tubulin code could steer nervous system physiology in vivo in general, and circuit development in particular, ^12^ is not well understood ^5, 13–15^.

Synaptic pruning—the wide-spread removal of exuberant synapses that is preserved across phyla, but also between development and disease—involves a local regulation of cytoskeletal function and stability ^16–19^. For instance, others and we have previously described the selective loss of microtubules in destabilized presynaptic axon branches ^16, 20, 21^. This loss can be mediated by the microtubule-severing enzyme, spastin ^20^, a protein commonly altered in hereditary spastic paraplegia ^8^. Indeed, microtubule destabilization, together with glia-mediated engulfment of synaptic material, is thus far one of very few shared mechanisms of synapse pruning across models and between development and disease ^22, 23^. The local loss of microtubules in pruning axon branches is accompanied by changes in PTMs ^20^, but whether these cytoskeletal changes are instructive or merely an epiphenomenon of the breakdown, and which molecular signals locally regulate microtubule modifications, remains unknown. Of particular significance in this context is polyglutamylation, a tubulin PTM that is highly enriched in the brain ^24^. Tubulin polyglutamylation involves the enzymatic “seeding” of an initial glutamate followed by addition of further glutamate residues, resulting in the formation of a polyglutamate (PolyE) chain. This PTM enhances the negative charge of the microtubule lattice, thus facilitating electrostatic interactions with neurodegeneration-associated MAPs, such as tau ^25, 26^, and severases, including spastin ^27^.

Based on these prior observations, we undertook a systematic investigation into the role of microtubule polyglutamylation during pruning. To this, we took advantage of the special features of synaptic pruning at the neuromuscular junction, a synapse which transitions within a few days from innervation by multiple to a single axon branch ^28^. Due to its accessibility and size, this synapse allows unique dynamic investigations with subsynaptic resolution ^20, 29–31^, which we used to reveal an instructive role of tubulin polyglutamylation in synaptic remodeling. Specifically, we show that in motor axon branches during pruning, polyglutamylation of tubulin alpha-4A (Tuba4a) results from the balanced action of the “writer” enzyme, tubulin tyrosine ligase-like glutamylase 1 (TTLL1; but not the functionally distinct TTLL7 ^13^) and the “eraser” enzymes, cytosolic carboxy peptidases (CCP) 1 and 6. The local density of polyglutamyl side chains on microtubules, modified in cell type-specific knock-out mice, in turn, determines the efficiency of spastin-mediated severing. We found that this interplay of PTMs and microtubule severing paces the speed by which peripheral synapses remodel, but confirmed a related mechanism for the timing of axon and spine pruning in the developing hippocampus.

## Results

### Glutamylases and deglutamylases are translationally regulated during motor axon pruning

To establish the expression patterns of polyglutamylation “writer” and “eraser” enzymes during motor axon pruning, we performed a translatome analysis of murine motor neurons using a ChAT-IRES-Cre X Rpl22^HA^ (called RiboTag hereafter) ^32, 33^ mouse cross-breeding. We immuno-isolated ribosome-bound mRNA from spinal cords, starting at postnatal day (P) 5 until 14 (P5, 7, 9, 11, 14; **Fig. 1a**). Specificity of this experiment for motor neuronal transcripts was verified by immunostaining for HA, which illustrated clear co-localization with Choline Acetyl Transferase (ChAT) (**Fig. 1b**), and quantitative PCR, which demonstrated a ≍20x enrichment of ChAT and absence of Glial Fibrillary Acidic Protein (GFAP) transcripts in the pull-downs compared to whole spinal cord mRNA (**Extended data, Fig. 1a** and **1b**). Principal component (PCA) analysis clearly separated the RiboTag pull-down vs. the total spinal cord mRNA samples (**Extended data, Fig. 1c**). In total, we identified > 4,500 differentially expressed transcripts at P9 (RiboTag vs whole spinal cord; Log_2_ |FC| ≥ 1.5; Padj ≤ 0.05; **Extended data, Table 1**). Gene ontology (GO) analysis of this gene list pointed towards neuron- or axon-specific biological processes (**Extended data, Fig. 1d** and **1e**). Similarly, gene set enrichment analysis revealed motor neuron markers (Isl1, Chat, Mnx1) augmented, while astrocyte markers (GFAP, Slc1a3, Aldh1l1, Olig2, Cspg4) and oligodendrocyte-specific genes (Sox10, Olig2, Cspg4) were depleted in the RiboTag pull-down samples (**Extended data, Fig. 1f**). We then scrutinized the complete gene list (**Extended data, Table 2**) obtained from our bona fide motor neuron translatome, for glutamylases (TTLL1, 3, 4, 5, 6, 7, 11 and 13) and deglutamylases (CCP1, 2, 3, 4, 5 and 6; **Extended data, Table 2**).

**Figure 1.**
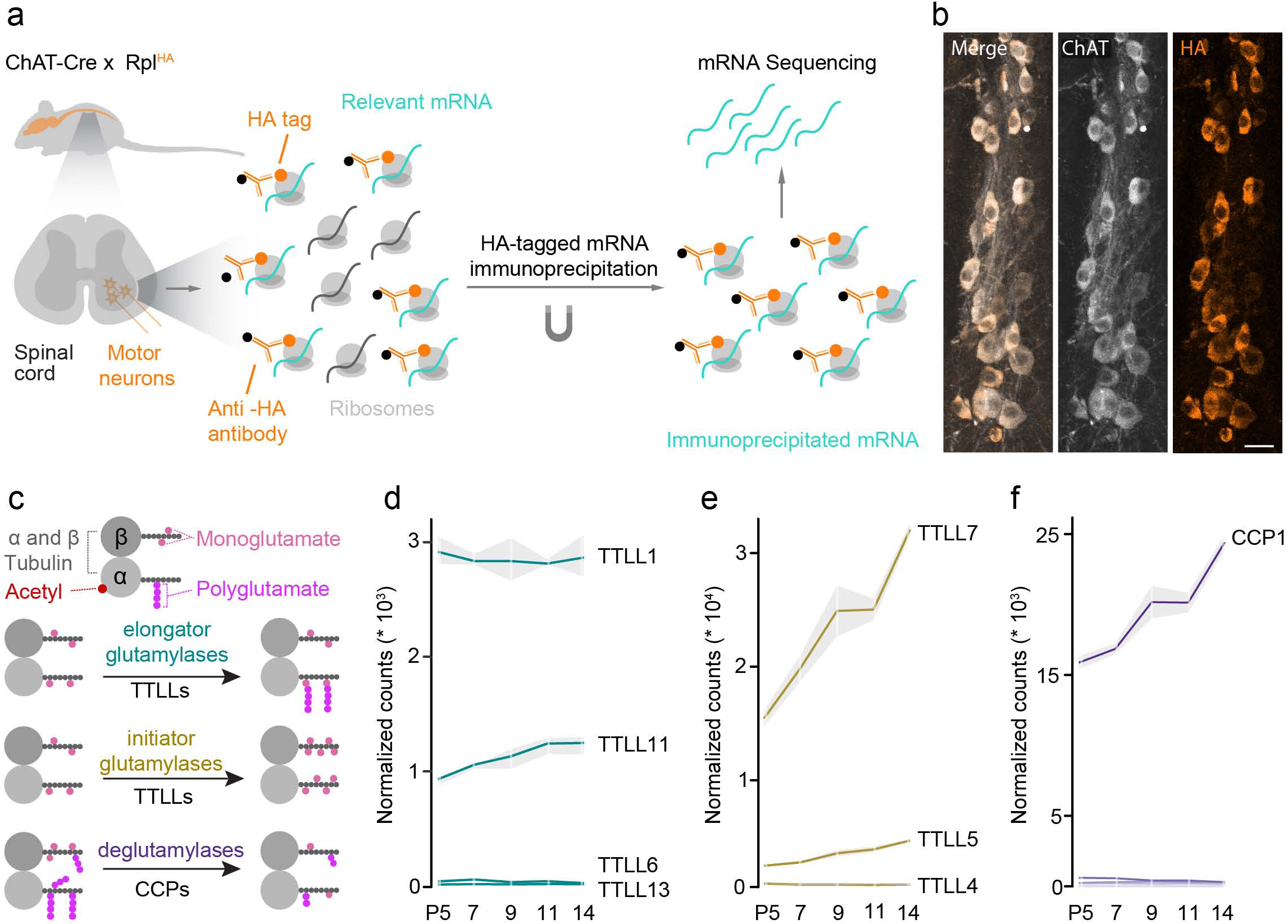
Motor neuron transcriptome during postnatal remodeling. **(a)** Strategy to isolate and sequence motor neuron specific mRNA from ChAT-Cre mice crossbred to Rpl22^HA^. HA-tagged ribosomes (orange) and associated mRNA (cyan) was immunoprecipitated from spinal cords at different developmental time points (postnatal day (P) 5, 7, 9, 11, 14) and sequenced. **(b)** Confocal image stacks of longitudinal spinal cord sections of 8-week-old ChAT-Cre X Rpl22^HA+^ mice immunostained for Choline acetyltransferase (ChAT, gray) and Hemagglutinin (HA, orange), merged channels on the left. Scale bar, 25 µm. **(c)** Schematic illustration of tubulin posttranslational modifications investigated in this study. Microtubules consist of tubulin alpha (α) and beta (β) dimers (gray), which may carry modifications in their luminal face (acetyl, red) or on their (-terminal tails (monoglutamate, pink; polyglutamate, magenta). Elongator TTLL glutamylases (turquoise) extend glutamyl side chains (polyglutamate, magenta); initiator TTLL glutamylases (gold) catalyze the addition of the first glutamate (monoglutamate, pink) to tubulin (-terminal tails; CCP deglutamylases (purple) remove glutamate residues. **(d) - (f)** Normalized mRNA read counts (*10^4^ or *10^3^) of **(d)** elongator TTLL glutamylases, **(e)** initiator TTLL glutamylases and **(f)** CCP deglutamylases in motor neurons at postnatal day PS, 7, 9, 11, 14. (n = 3 animals per age group). Graphs: mean ± SEM.

“Writer” TTLL glutamylases decorate the microtubule lattice with glutamate either in a “seeder” fashion, by preferentially adding the first branch glutamate (E) onto the tubulin C-terminal tail (monoE), or by elongating previously seeded E-chains ^13, 34^ (**Fig. 1c**). Among the known “elongators”, TTLL1 showed the highest expression levels (e.g., at P14: approximately 69% of “elongator” TTLL levels) that were stable across all analyzed time points. In contrast, TTLL11 was expressed at a substantially lower level (e.g., at P14: 29 %), while TTLL6 and TTLL13 were virtually absent (**Fig. 1d**). Analysis of “seeders” yielded increasing TTLL7 mRNA reads between P5 and P14 (e.g., at P14: 87 % of “seeder” TTLL levels), while TTLL5 and TTLL4 transcripts were much less abundant (at P14: 11 % and 1 % for TTLL5 and TTLL4, respectively; **Fig. 1e**). Amongst the deglutamylases, CCP1 transcripts were the most abundant and increased during the pruning phases (e.g., increase from P5 to P14: 53 %; **Fig. 1f**). Among all the other CCP-type deglutamylases (CCP2, 3, 4, 5, 6), CCP6 was the most abundant enzyme which exhibited an increase of 28% between P5 and P14. The rest were present at very low transcript rates, ranging e.g. from 0.03% to 1.1% at P14 (**Fig. 1f**). These findings are consistent with previous reports showing that CCP1 and CCP6 are the two most abundant deglutamylases in the central nervous system ^35^. Thus, we focused our efforts on genetic disruption of TTLL1, TTLL7, CCP1 and CCP6 to probe their role as potential candidates that mediate polyE-instructed neuronal remodeling. We first generated cholinergic neuron-conditional mouse mutants of the most expressed glutamylases, i.e. TTLL1^mnKO^ (ChAT-IRES-Cre X TTLL1^flox/flox^; see experimental procedures), which should mostly affect polyglutamylation on tubulin alpha ^13^, and TTLL7^mnKO^ (ChAT-IRES-Cre X TTLL7^flox/flox^; see experimental procedures), which would be expected to deplete monoglutamylation on tubulin beta ^13^.

### Genetic ablation of TTLL1, but not of TTLL7, delays neuromuscular synapse elimination

We quantified microtubule content and PTM patterns by immunohistochemistry and confocal microscopy in the triangularis sterni muscles of these mice at P9 ^20, 31, 36^. To determine microtubular content, we used an anti-tubulin beta-3 (Tubb3) antibody, the most abundant tubulin beta isoform in motor neurons after P9 (**Extended data, Fig. 1g**), while an anti-PolyE antibody, which recognizes C-terminal chains of ≥ 3 glutamate residues on tubulin tails, was used to assess axonal polyglutamylation levels. As expected, PolyE normalized to Tubb3 was drastically reduced (to 0.09-fold), while microtubule content was 2.8-fold increased (normalized to neurofilament; **Fig. 2a**-**d**). In contrast, immunostaining using an anti-βmonoE antibody (normalized to Tubb3), which detects a single glutamate seed on E435 of tubulin beta-2 tails, was largely unaffected (**Fig. 2e** and **2f**). Finally, antibody staining for acetylated tubulin (acetyl-K40) was also decreased (0.68-fold, normalized to Tubb3, **Fig. 2g** and **2h**), in line with previous reports ^37^. Thus, microtubules in motor axons in TTLL1^mnKO^ mice are more abundant but may carry less PTMs. As revealed by time-lapse imaging of nerve-muscle-explants of TTLL1^mnKO^ that also carried a Thy1-EB3-YFP transgene ^38^, most parameters of plus-end polymerization dynamics were unaffected (including comet density, orientation and velocity, **Extended data, Fig. 2a, 2c and 2d**), but comet length distribution was significantly, albeit mildly altered in TTLL1^mnKO^ compared to TTLL1^mnWT^ (**Extended data, Fig. 2b**). Specifically, EB3 comet elongation in TTLL1^mnKO^ increased aligning with recent in vitro findings that show augmented microtubule growth rate upon deglutamylation ^39^. Concomitant to the increased microtubule mass and reduced PolyE levels, a higher percentage of neuromuscular synapses remained innervated by two or more axons in TTLL1^mnKO^ vs. control muscles at all postnatal ages tested (**Fig. 2i** and **2j**). Collectively these findings suggest that TTLL1-mediated polyglutamylation instructs the local dismantling of microtubules, and thus paces developmental axon pruning of motor neurons.

**Figure 2.**
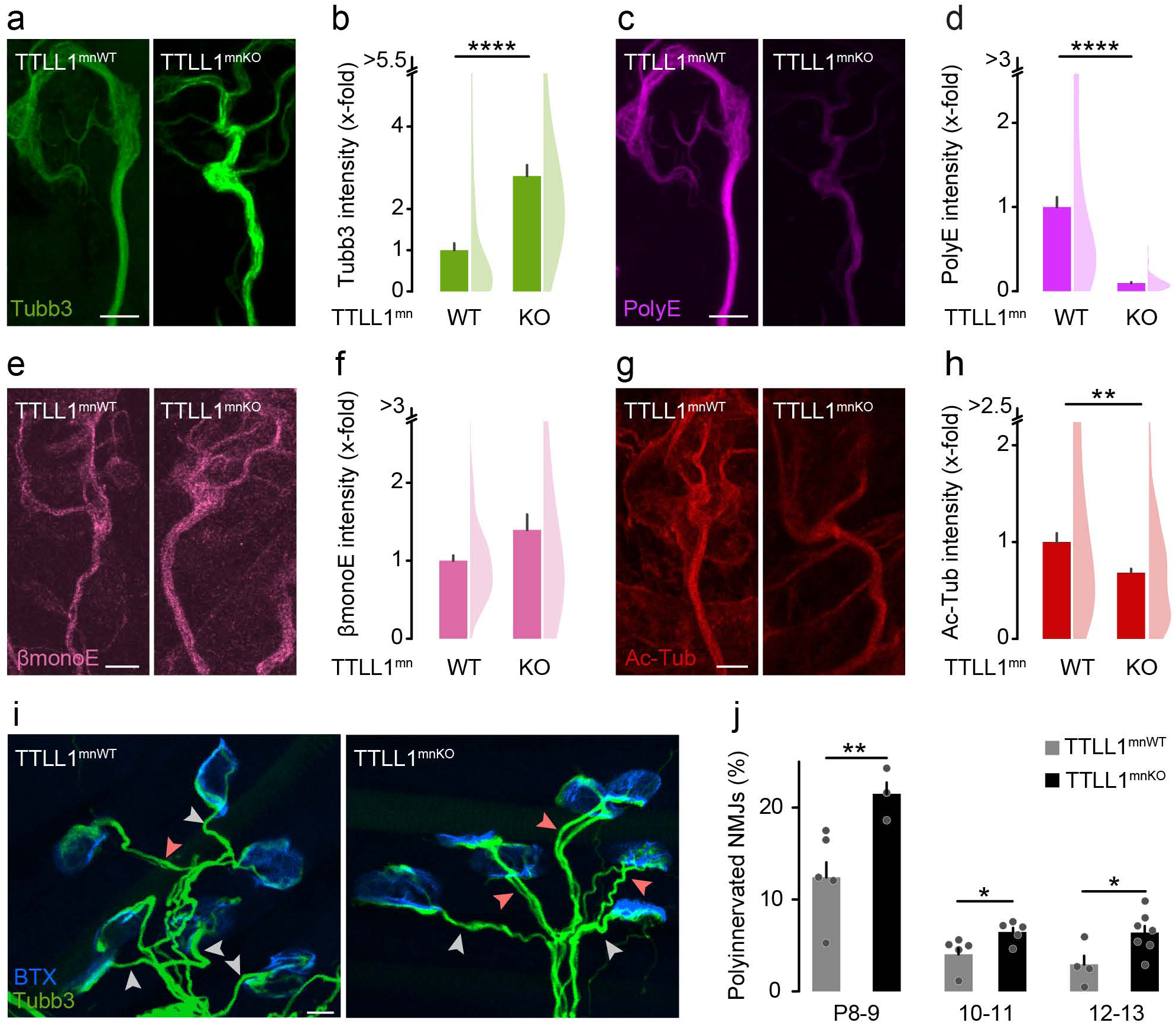
Genetic ablation of TTLLl delays motor axon remodeling. **(a) - (h)** Quantitative immunostainings for microtubule markers and neurofilament heavy polypeptide on triangularis sterni muscles at P8-9 from TTLLl^mnKO^ and TTLLl^mnWT^ littermate controls. **(a)** Confocal stack of a neuromuscular synapse depicting tubulin beta-3 staining (Tubb3, green). **(b)** Quantification of Tubb3 intensity in terminal motor axons, normalized to neurofilament heavy polypetide (n = 3 mice per genotype, n 70 axons per genotype). **(c)** Confocal stack of a neuromuscular synapse depicting polyglutamate chain staining (PolyE, magenta). **(d)** Quantification of polyE intensity in terminal motor axons, normalized to tubulin beta-3 (n = 3 mice per genotype, n 70 axons per genotype). **(e)** Confocal stack of a neuromuscular synapse depicting monoglutamate on tubulin beta isoform staining (monoE, pink). **(f)** Quantification of monoE intensity in terminal motor axons, normalized to tubulin beta-3 (n = 3 mice per genotype, n 54 axons per genotype). **(g)** Confocal stack of a neuromuscular synapse depicting acetylated tubulin staining (Ac-Tub, red). **(h)** Quantification of ac-tub intensity in terminal motor axons, normalized to tubulin beta-3 (n 4 mice per genotype, n 68 axons per genotype). **(i)** Confocal stacks of neuromuscular synapses of P12 TTLLl^mnKO^ and TTLLl^mnWT^ littermates crossbred to Thyl-YFP (Tubb3, green; BTX, blue), depicting singly (gray arrowheads) and polyinnervated synapses (light red arrowheads). **(j)** Percentage of polyinnervated neuromuscular junctions (NMJs) in TTLL1mnWT vs TTLL1mnKO littermates crossbred to Thyl-YFP at P8-9, 10-11, 12-13. (n 3 mice per group, n 197 NMJs per animal). Each dot represents one animal. Graphs: mean+ SEM (left) and data representing axons as half violin (right) (b,d,f,h) or mean+ SEM and data representing animals as single dots (j). Mann-Whitney test determined significance:*, P < 0.05; **, P< 0.01; ****, P < 0.0001. Scale bars (a, c, e, g) 5 µm, (i) 10 µm.

In contrast, the same set of analyses in TTLL7^mnKO^ mice revealed no changes in overall microtubule mass and dynamism in terminal motor axons (**Fig. 3a** and **3b**; **Extended data, Fig. 2e-2h**). In line with the specificity of TTLL7 activity ^13^, the intensity of PolyE immunostaining was not altered (**Fig. 3c** and **3d**), while quantification of βmonoE immunofluorescence revealed a decrease compared to control axons as expected (0.55-fold, normalized to Tubb3, **Fig. 3e** and **3f**). Consistently, the pruning speed of motor axons was similar in TTLL7^mnKO^ vs. control muscles (**Fig. 3g**). These data corroborate that only the density of TTLL1-added polyglutamylation, but not TTLL7-catalyzed βmonoE seeds, influences pruning—suggesting a high level of specificity in the regulation of pruning via microtubule PTMs, in line with the tubulin code concept.

**Figure 3.**
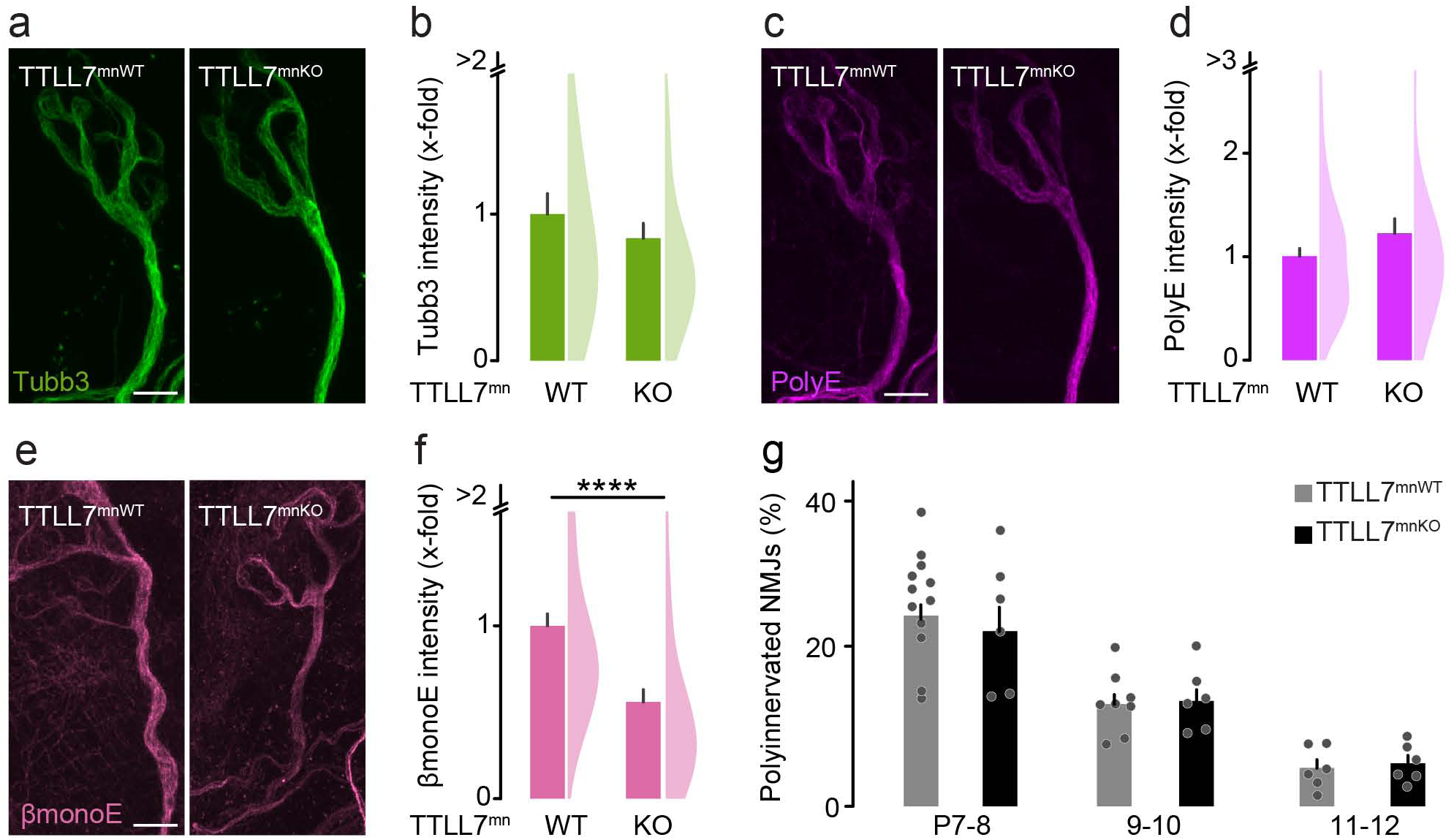
Genetic ablation of TTLL7 does not affect peripheral pruning. **(a) - (f)** Quantitative immunostainings for microtubule markers and neurofilament heavy polypeptide on triangularis sterni muscles at pg from TTLL7^mnKO^ and TTLL7^mnWT^ littermate controls. **(a)** Confocal stack of a neuromuscular synapse depicting tubulin beta-3 staining (Tubb3, green). **(b)** Quantification of Tubb3 intensity in terminal motor axons, normalized to neurofilament heavy polypetide (n 4 mice per group, n 55 axons per animal). **(c)** Confocal stack of a neuromuscular synapse depicting polyglutamate chain staining (PolyE, magenta). **(d)** Quantification of polyE intensity in terminal motor axons, normalized to tubulin beta-3 (n 4 mice per group, n 55 axons per animal). **(e)** Confocal stack of a neuromuscular synapse depicting monoglutamate on tubulin beta isoform staining (monoE, pink). **(f)** Quantification of monoE intensity in terminal motor axons, normalized to tubulin beta-3 (n 4 mice per genotype, n 76 axons per genotype). **(g)** Percentage of polyinnervated neuromuscular junctions (NMJs) in TTLL7^mnWT^ vs TTLL7^mnKO^ littermates crossbred to Thyl-YFP at P7-8, 9-10, and 11-12. (n 6 mice per group, n 96 NMJs per animal). Graphs: mean+ SEM (left) and data representing axons as half violin (right) (b,d,f) or mean+ SEM and data representing animals as single dots (g). Mann-Whitney test determined significance: ****, P < 0.0001. Scale bars, 5 µm.

### TTLL1^KO^ leads to pruning defects in the CNS

We next explored whether reduced TTLL1-mediated polyglutamylation of microtubules would also result in defective pruning processes in the CNS. We chose granule neurons of the dentate gyrus, which provide a well-established and easily quantified murine model for ‘stereotyped’ pruning. During the first two postnatal months, granule cells retract their infrapyramidal bundle (IPB) of mossy fibers, while the main suprapyramidal bundle (SPB), is maintained (see schematic; **Fig. 4a**). We used a constitutive TTLL1^KO^ mouse model, which exhibits reduced PolyE levels in the brain ^14^. Quantification of the IPB/SPB length ratio in sections immunostained for Calbindin D-28K, which selectively stains mossy fibers, revealed delayed pruning of the IPB in TTLL1^KO^ vs. control mice (**Fig. 4b-d**). We further tested if elimination of spines in granule neuron dendrites —i.e., remodeling of input synapses further upstream in hippocampal circuitry—was delayed as well. Gene-gun based DiI labelling ^40^ before (3 week-old mice) and after the pruning phase (8 week-old mice) showed that TTLL1^KO^ mice developed normal spine densities initially, but failed to prune (**Fig. 4e-g**). This suggests that TTLL1-mediated polyglutamylation might be a general pacemaker for the execution of remodeling processes, not only during competitive elimination of peripheral synapses, but also during stereotyped pruning in the CNS.

**Figure 4.**
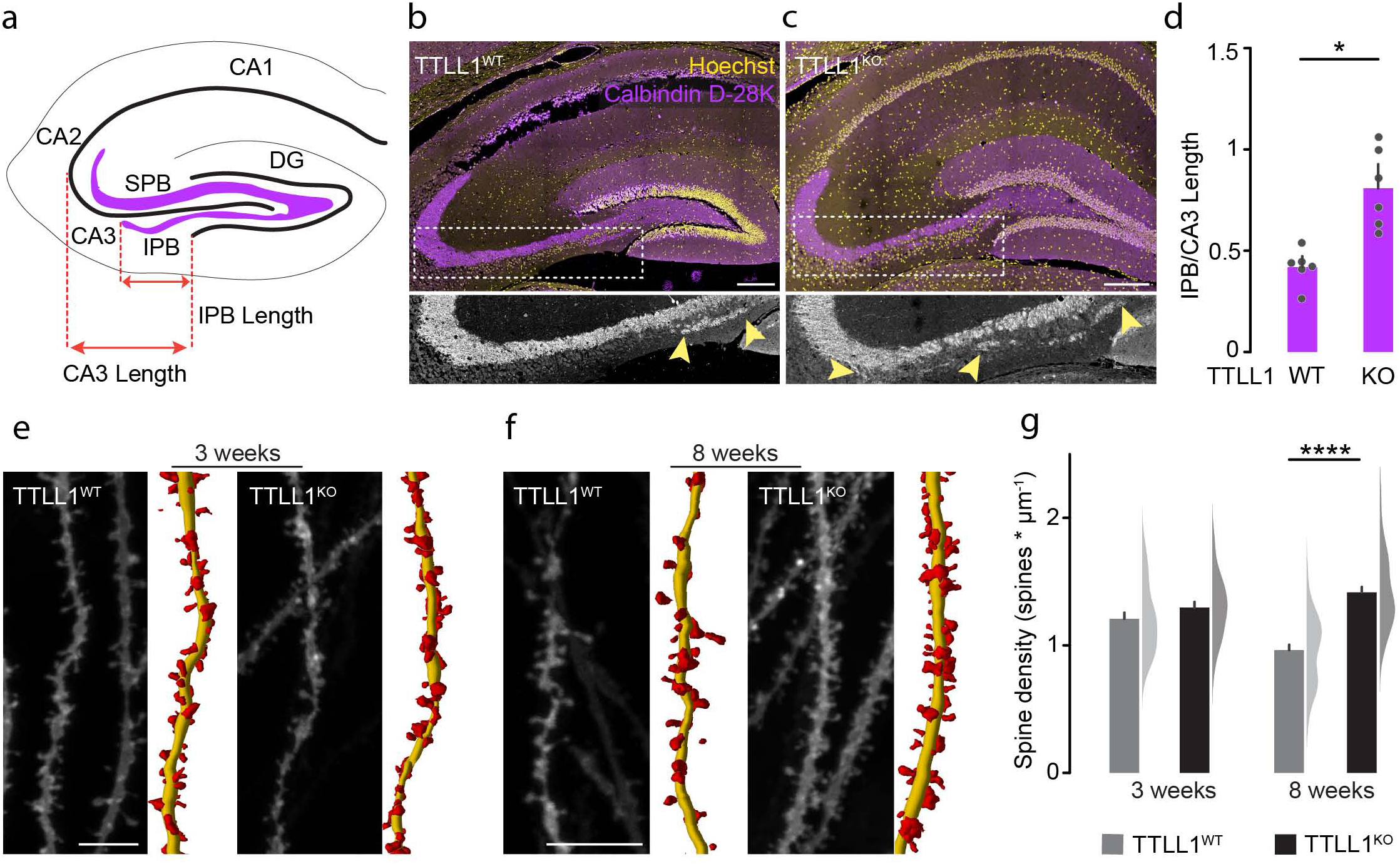
TTL1^KO^ leads to pruning defects in the CNS. **(a)** Schematic of the hippocampus showing infrapyramidal bundle (IPB, violet), suprapyramidal main bundle (SPB, violet), dentate gyrus (DG), CAl, CA2, CA3 region; red arrows indicate IPB and CA3 length used in quantification in (d). **(b)-(c)** Confocal stacks of coronal brain sections of 8-week-old TTLLl^WT^ vs TTLLl^KO^ littermates immunostained for Calbindin D-28k (violet) and Hoechst (yellow). Boxed area shows examples of IPB length measurement (yellow arrowheads). **(d)** Quantification of IPB length normalized to CA3 in 8-week-old TTLLl^WT^ vs TTLLl^KO^ littermates (n=6 animals per genotype). **(e)** - **(f)** Confocal stacks of brain slices of TTLLl^WT^ and TTLLl^KO^ mice depicting Dil-labelled dendritic spines of hippocampal granule cells, at **(e)** 3 and **(f)** 8 weeks of age. Right side panels illustrate spine reconstructions (dendrites, yellow; spines, red). **(g)** Quantification of dendritic spine density in TTLLl^WT^ vs TTLLl^KO^ at 3 weeks (n 2: 30 spines from 6 animals) and 8 weeks (n 2: 43 spines from 8 animals). Graphs: mean+ SEM and data representing animals as single dots (d) or mean+ SEM (left) and data representing dendritic spines as half violin (right) (g). Mann-Whitney test determined significance.*, P < 0.05; ****, P < 0.0001. Scale bars, 100 µm (b, c) and 10 µm (e, f).

### Polyglutamylation of tubulin alpha-4A instructs remodeling

Exploration of tubulin isoforms in our motor neuron translatome data revealed an upregulation of tubulin alpha-4A (Tuba4a) among the alpha isoforms, with a consistent increase across the synaptic pruning phase (+63% from P5 to P14; **Fig. 5a**). In contrast, other tubulin alpha isoforms were either stably expressed (e.g., Tuba1b and Tuba1c; **Fig. 5a**) or downregulated (Tuba1a; **Fig. 5a**). Interestingly, Tuba4a is the isoform which seems to carry the longest glutamate side chains ^41, 42^. In light of this expression pattern, we investigated the effect of Tuba4a-specific polyglutamylation loss in neuronal remodeling. For this, we utilized Tuba4a knock-in mice (Tuba4a^KI^), which carry a mutated Tuba4a allele, which prohibits glutamylation of this specific tubulin isoform ^41^. Using confocal analysis of double immunostainings for PolyE and Tubb3 in terminal axons in Tuba4a^KI^ triangularis sterni muscles, we found a phenotype resembling to that of TTLL1^mnKO^: Tubb3 intensity increased (1.2-fold; normalized to neurofilament; **Fig. 5b** and **5c**), while the intensity of the PolyE signal staining was reduced (0.9-fold; normalized to Tubb3; **Fig. 5d** and **5e**). As motor axon remodeling was significantly, albeit transiently, delayed (**Fig. 5f**), we conclude that polyglutamylation of tubulin alpha-4A is necessary to initially drive the pruning of motor axons.

**Figure 5.**
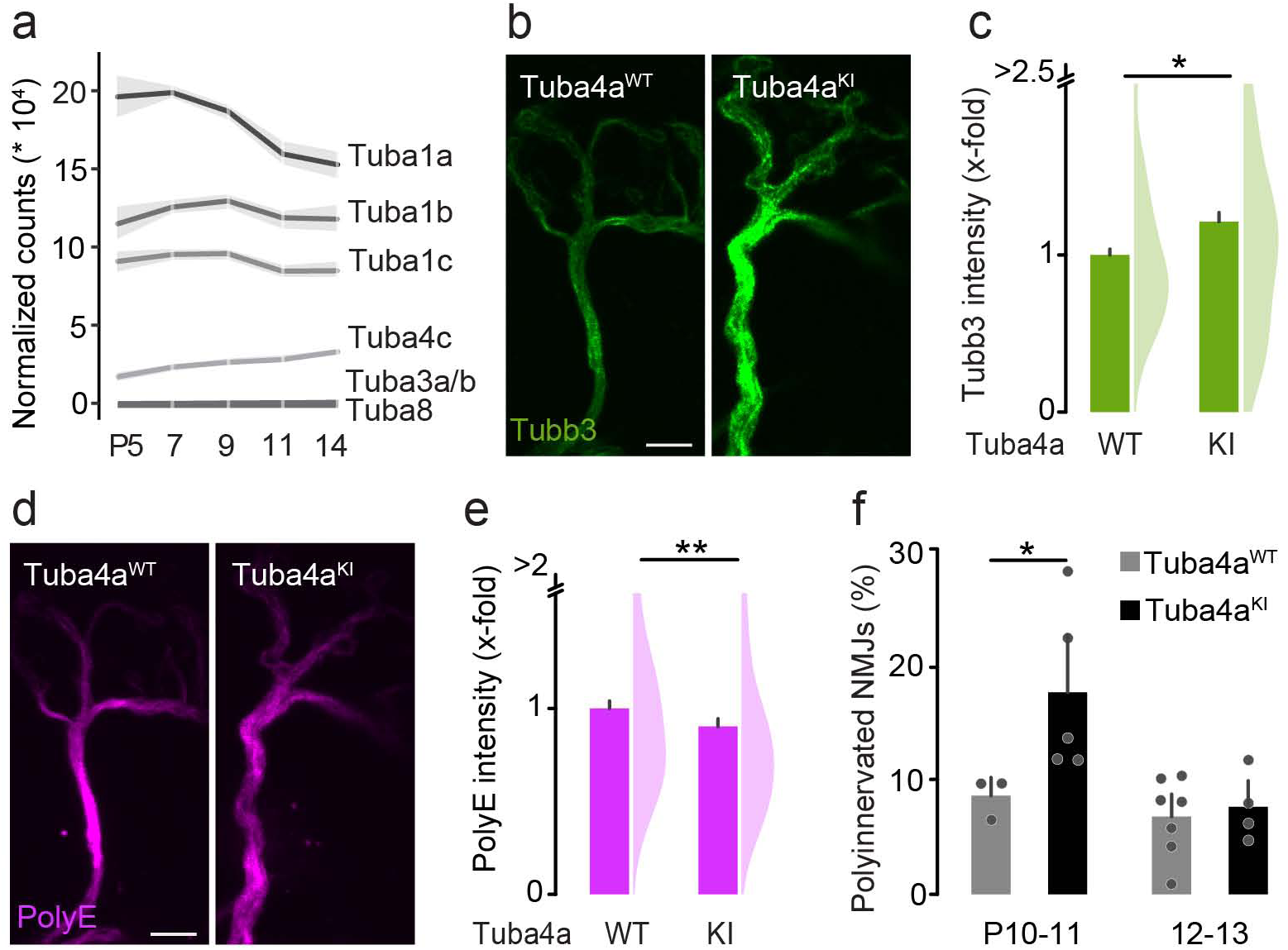
Polyglutamylation of tubulina4A instructs remodeling. **(a)** Graph depicts normalized mRNA counts (*10^4^) oftubulin alpha isoforms across the motor axon remodeling phase (postnatal day (P) 5, 7, 9, 11, 14). **(b)-(e)** Quantitative immunostainings for tubulin beta-3, neurofilament heavy polypeptide and polyglutamate on triangularis sterni muscles at Pl0-11 and 12-13 from Tuba4a^KI^ and Tuba4a^WT^ littermates. **(b)** Confocal stack of a neuromuscular synapse depicting tubulin beta-3 staining (Tubb3, green). **(c)** Quantification of Tubb3 intensity in terminal motor axons, normalized to neurofilament heavy polypetide (n 10 mice per genotype, n 174 axons per animal). **(d)** Confocal stack of a neuromuscular synapse depicting polyglutamate chain staining (PolyE, magenta). **(e)** Quantification of polyE intensity in terminal motor axons, normalized to tubulin beta-3 (n 10 mice per genotype, n 174 axons per animal). **(f)** Percentage of polyinnervated neuromuscular junctions (NMJs) in Tuba4a^WT^ vs Tuba4aK^1^littermate controls, based on tubulin beta-3 staining, at Pl0-11 and 12-13. (n 3 mice per group, n 60 NMJs per animal) Graphs: mean± SEM (a), mean + SEM (left) and data representing axons as half violin (right) (c,e) or mean+ SEM and data representing animals as single dots (f). Mann-Whitney test determined significance: *, P < 0.05; ****, P < 0.0001. Scale bars, 5 µm.

### Genetic deletion of deglutamylases CCP1 and CCP6 in motor neurons accelerates pruning

We next wondered whether the described system operates in both directions, meaning that hyper-polyglutamylation of microtubules would accelerate axon pruning. Deglutamylases (CCPs) catalyze the removal of glutamate chains from tubulin C-terminal tails ^43–45^, and thus counteract the enzymatic activity of elongating glutamylases. We focused our efforts on combined motor neuron-specific deletions of CCP1 and CCP6 (CCP1&6^mnKO^), as—while CCP1 was the dominant CCP transcript in our translatome data (**Fig. 1f**)—CCP6 is known to have compensatory potential.

Immunofluorescence analysis in terminal triangularis sterni axons at P9 revealed a reduction of Tubb3 levels in CCP1&6^mnKO^ mice compared to littermate controls (0.8-fold; normalized to neurofilament; **Fig. 6a** and **6b**). Unexpectedly, PolyE immunostaining intensity was significantly decreased (0.6-fold; normalized to Tubb3; **Fig. 6c** and **6d**), while βmonoE levels remained unaltered as predicted (**Fig. 6e** and **6f**). In converse to the TTLL1^mnKO^ phenotype, acetylated microtubules were increased significantly (**Fig. 6g** and **6h**). Furthermore, we observed changes of EB3-YFP comet dynamics, with a 27 % drop in density and a 12 % increase in mean velocity in CCP1&6^mnKO^ compared to control mice(**Extended data, Fig. 2i** and **2k**). Microtubule plus-end growth (EB3 comet orientation) and length distribution remained normal (**Extended data, Fig. 2j and 2l**). Correlative live imaging of EB3-YFP comet density normalized to tubulin beta-3 immunostaining in fixed tissue on the single axon level was unchanged in the CCP1&6^mnKO^ compared to CCP1&6^mnWT^ controls (**Extended data, Fig. 3a-d**). Of note, the synapse elimination process in CCP1&6^mnKO^ mice was accelerated, in accordance with lower Tubb3 levels, as fewer neuromuscular synapses were polyinnervated at several time points (**Fig. 6i**). This finding corroborates that the polyglutamylation “writer” and “eraser” enzymes bidirectionally or “rheostatically” regulate the developmental remodeling of axons and synapses.

**Figure 6.**
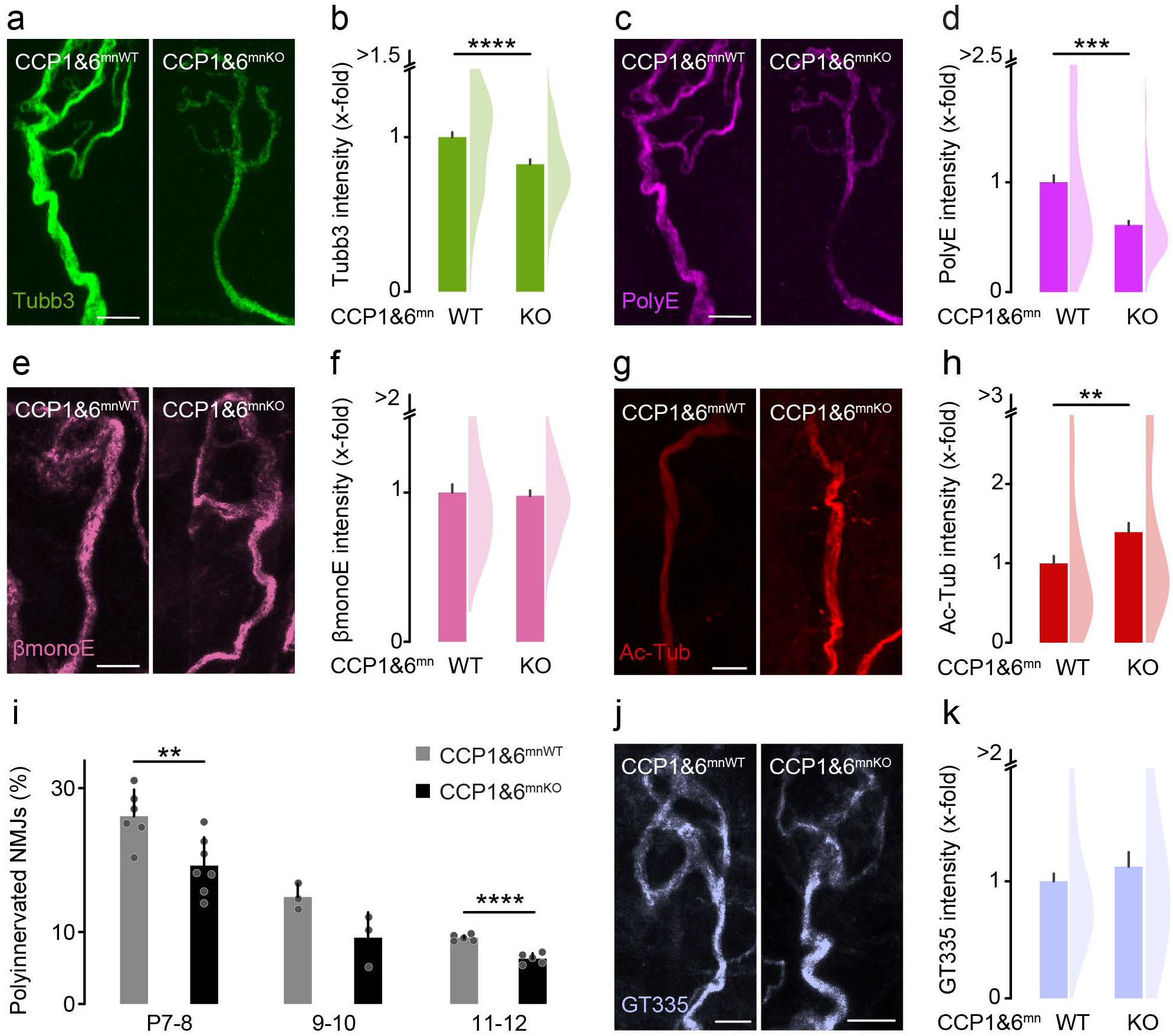
Genetic deletion of CCPl and CCP6 in motor neurons accelerates pruning. **(a) - (j)** Quantitative immunostainings for microtubule markers and neurofilament heavy polypeptide on triangularis sterni muscles at pg from CCP1&6^mnKO^ and CCP1&6^mnWT^ littermate controls. **(a)** Confocal stack of a neuromuscular synapse depicting tubulin beta-3 staining (Tubb3, green). **(b)** Quantification of Tubb3 intensity in terminal motor axons, normalized to neurofilament heavy polypetide (n ≥ mice per genotype, n ≥ 74 axons per genotype). **(c)** Confocal stack of a neuromuscular synapse depicting polyglutamate chain staining (PolyE, magenta). **(d)** Quantification of polyE intensity in terminal motor axons, normalized to tubulin beta-3 (n ≥ mice per genotype, n 74 axons per genotype). **(e)** Confocal stack of a neuromuscular synapse depicting monoglutamate on tubulin beta isoform staining (βmonoE, pink). **(f)** Quantification of monoE intensity in terminal motor axons, normalized to tubulin beta-3 (n ≥ 3 mice per genotype, n ≥ 63 axons per genotype). **(g)** Confocal stack of a neuromuscular synapse depicting acetylated tubulin staining (Ac-Tub, red). **(h)** Quantification of ac-tub intensity in terminal motor axons, normalized to tubulin beta-3 (n ≥ 4 mice per genotype, n ≥ 80 axons per genotype). (i) Percentage of polyinnervated neuromuscular junctions (NMJs) in CCP1&6^mnWT^ vs CCP1&6^mnKO^ littermates crossbred to Thyl-YFP across at P7-8, 9-10, and 11-12. (n ≥ 3 mice per group, n ≥ 125 NMJs per animal). **(j)** Confocal stack of a neuromuscular synapse depicting staining of branching points of glutamate side chains on tubulin beta and alpha (GT335, light blue). **(k)** Quantification of GT335 intensity in terminal motor axons, normalized to tubulin beta-3 (n ≥ 4 mice per genotype, n 90 axons per genotype). Graphs: mean+ SEM (left) and data representing axons as half violin (right) (b,d,f,h,j) or mean+ SEM and data representing animals as single dots (k). Mann-Whitney test determined significance:**, P < 0.01; ***, P < 0.001; ****, P < 0.0001. Scale bars 5 µm.

### Spastin swiftly eliminates excessively polyglutamylated microtubules

The absence of excess polyglutamylation in young CCP1&6^mnKO^ motor axons is surprising, as CCPs remove glutamate residues from microtubules; consequently, their genetic deletion should result in an increased PolyE immunostaining. We explored two possible explanations for this observation: (i) excessive polyglutamylation might result in highly branched, arborized polyglutamyl chains that could evade PolyE-immunodetection due to steric hindrance ^46^ or (ii) instantaneous severing and subsequent destabilization of excessively polyglutamyl-decorated microtubules by enzymes such as spastin ^27^ might leave the polyglutamylated microtubule pool paradoxically depleted.

To test the former possibility, we performed anti-GT335 immunostainings, an antibody recognizing the branching point of glutamate chains on tubulin tails ^24, 47^ independent of chain length. Confocal quantification indicated a similar staining intensity in CCP1&6^mnKO^ mice compared to controls (**Fig. 6j** and **6k**). This suggests that the absence of hyper-polyglutamylation in CCP1&6^mnKO^ mice has a biological, rather than technical, explanation. Next, we focused on a possible intersection of the effects of absent CCP-type deglutamylase activity and spastin activation. To test this, we conducted neonatal (P3) injections of an adeno associated virus (AAV9-hSyn-Cre), which expressed Cre recombinase in neurons only, in conditional CCP1 and spastin mice that also carried a reporter allele (CCP1^flox/flox^ X Spast^flox/flox^ X TdTomato). The TdTomato reporter fluorescence was analyzed in motor axons at P9 to validate successful Cre-mediated excision (**Fig. 7a**). To circumvent background ambiguities and the lack of reliable in situ detection tools for endogenous spastin or deglutamylases, the tdTomato signal was indexed in quantiles based on fluorescent intensity. We compared the quantile with the highest intensity (tdTomato^high^; which we considered most likely for CCP1 and spastin double-deletion) with the bottom quantile (tdTomato^low^; likely non-recombined) for levels of PolyE immunostaining. This indeed revealed a 2-fold increase of PolyE in the tdTomato^high^ compared to the tdTomato^low^ axons (**Fig. 7b** and **7d**), which was accompanied by added microtubule content (1.3-fold; **Fig. 7b** and **7c**). Notably, spastin translation decreases from P5 onwards (which contrasts to other severing enzymes, such as Katnal1; **Extended data, Fig. 4a** and **Table 3**). This suggests that deglutamylases might be more relevant later in life—and, consistently, quantification of immunostainings in adult CCP1&6^mnKO^ motor axons showed lower microtubule mass (0.4-fold; normalized to neurofilament; **Extended data, Fig. 4b** and **4c**) accompanied by a drastic increase in PolyE intensity (4.1-fold; **Extended data, Fig. 4d** and **4e**). In summary, microtubular content and PTM composition in motor axons appears to be controlled by the interplay of tubulin glutamylases, deglutamylases and specific microtubule severing enzymes.

**Figure 7.**
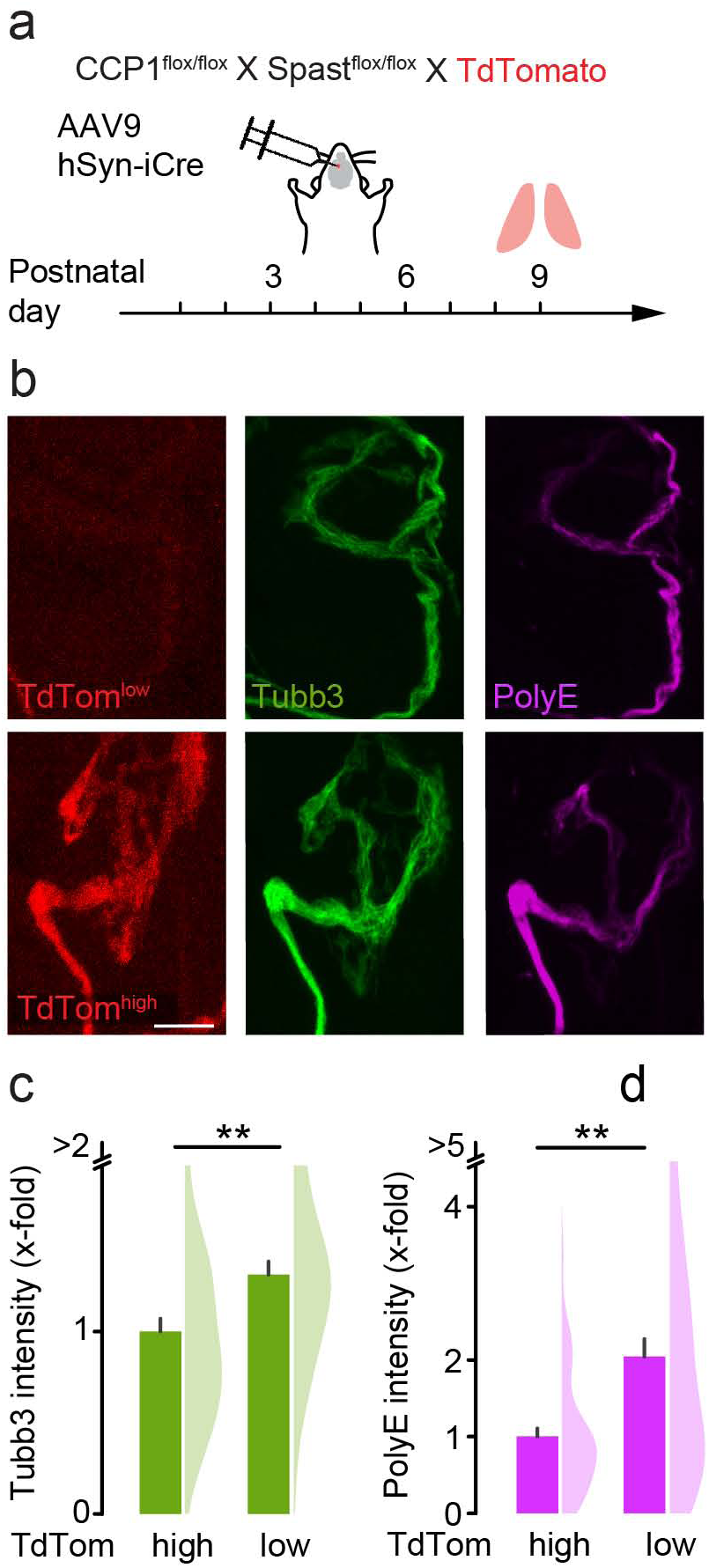
Modulation of polyglutamylation by deletion of spastin and CCPl. **(a)** Schematic for acute deletion of CCPl and spastin in a subset of motor neurons. P3 CCPl^flox/flox^ X Spastl^flox/flox^ X TdTomato reporter animals were intraventricularly injected with viral vectors (AAV9-hSyn-iCre) and immunostained for microtubule markers at P9. Expression of TdTomato (TdTom) served as a measure for CCPl and Spastin deletion. Terminal motor axons were considered Cre recombined (TdTom^high^) if their TdTomato intensity belonged to the 3rd quantile range, and not recombined if they belonged to 1st quantile range (TdTom^low^). **(b)** Confocal stacks of neuromuscular synapses depicting TdTomato signal and tubulin beta-3 and polyglutamate staining (TdTomato, red; Tubb3, green; PolyE, magenta). **(c)** Quantification of Tubb3 intensity in terminal motor axons (n 6 mice per group, n 50 axons per group). **(d)** Quantification of polyE intensity in terminal motor axons, normalized to Tubb3 (n 6 mice, n 50 axons per group). Graphs: mean + SEM (left) and data representing axons as half violin (right) (c,d). Mann-Whitney test determined significance: **, P < 0.01; ***, P < 0.001. Scale bars 5 µm.

### Microtubular PTM levels are under control of neurotransmission

Neuronal activity is a main driver of sculpting neural circuits, including at the neuromuscular synapse ^19^, and it has been demonstrated to facilitate tubulin polyglutamylation ^48^. We previously demonstrated that block of neurotransmission by injection of α-bungarotoxin (α-BTX), which blocks postsynaptic acetylcholine receptors and stalls synapse elimination, leads to a reduction of tubulin beta-3 immunofluorescence in the presynaptic motor axon ^49^. In order to test if this loss of tubulin beta-3 is mediated by activity-driven modulation of PTMs, we analyzed polyglutamylation in motor axons at P9 following an injection of α-BTX into the thorax two days earlier (**Fig. 8a**). In axon branches that terminated in α-BTX-blocked synapses (BTX+) polyglutamylation dropped significantly compared to control axons (0.7-fold; normalized to Tubb3; **Fig. 8b** and **8c**). This reduction following block of neurotransmission is reminiscent of CCP1&6^mnKO^ mice, where spastin appears to deplete polyglutamylated microtubules faster than glutamyl side chains can accumulate. In order to test this notion, we injected α-BTX into mice that globally lack spastin (Spast^KO^) and accumulate polyglutamylation ^20^, enabling us to detect glutamylase activity upon block of neurotransmission. Indeed, PolyE immunostaining levels were increased (2.1-fold) in blocked axons (α-BTX+/Spast^KO^) compared to branches innervating unaffected synapses (α-BTX-/Spast^KO^; **Fig. 8b** and **8c**). Thus, our data suggest that the enzymatic activity of the “elongator” TTLL glutamylase in the presynaptic axons is governed by neurotransmission. In summary, polyglutamylation indeed appears to “encode” activity-dependent signals that are subsequently “read” by spastin, which severs polyglutamylated microtubules and thus facilitates axon branch and synapse removal.

**Figure 8.**
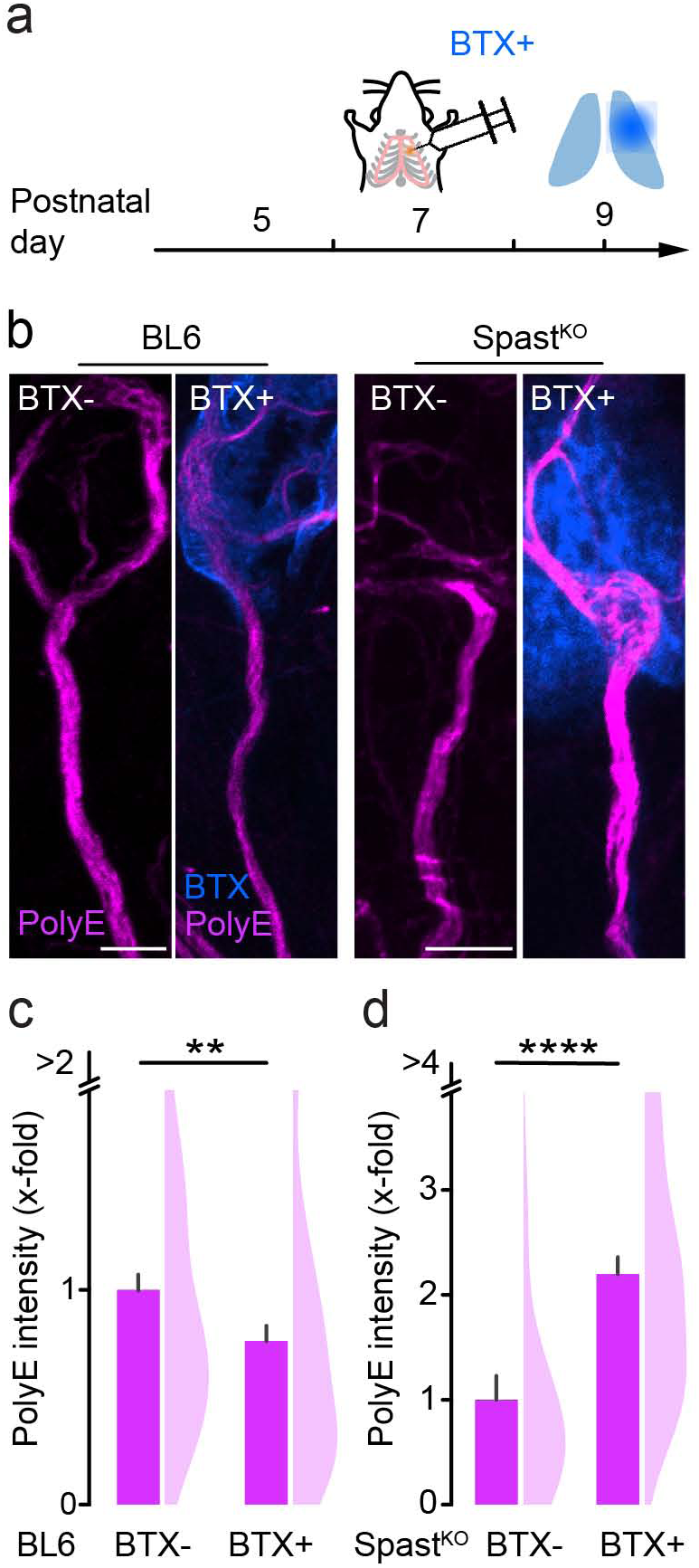
Modulation of polyglutamylation by blockage of neurotransmission. **(a)** Schematic for injections of α-bungarotoxin (BTX) conjugated to Alexa Fluor 594. C57BL/6N (BL6) and Spast^KO^ mice were unilaterally injected with α-BTX into the thoracic wall at P7. Muscles were fixed and immunostained at P9 for microtubule markers. **(b)** Confocal stacks of neuromuscular synapses, depicting non-injected motor axons stained for polyglutamate and BTX-injected motor axons (BTX, blue) stained for polyglutamate (PolyE, magenta), at pg in BL6 (left) and Spast^KO^ (right) animals. **(c)** - **(d)** Quantification of polyE intensity normalized to Tubb3 in terminal motor axons of **(c)** BL6 mice (n = 4 mice per group, n 87 axons per group) and **(d)** Spast^KO^ mice (n 1 mice per group, n 23 axons per group). Graphs: mean + SEM (left) and data representing axons as half violin (right). Mann-Whitney determined significance: *, P < 0.05; ***, P < 0.001; ****, P < 0.0001. Scale bars, 5 µm.

## Discussion

Neurons remodel extensively during development, with remarkable temporal and spatial precision ^19,50^. The fate of adjacent and otherwise indistinguishable neurites and synapses can starkly diverge— some are swiftly removed while others, often immediate neighbors, last for a lifetime ^51^. The microtubule cytoskeleton appears to play a key role in such remodeling processes ^16, 31^. Albeit, how cytoskeletal functions would be regulated in the required temporally- and spatially-controlled manner has remained unclear. Here, we reveal a rheostatic system of enzymes ^27^ that determines the speed of remodeling and is governed by the “writer” polyglutamylase TTLL1 and opposing “eraser” CCP deglutamylases. This system sets the glutamylation level, which in turn determines axon or synapse fate via the microtubule destabilizing action of the “reader” severase spastin ^20^.

This concept is noteworthy from two vantage points: First, as a unique in vivo manifestation of the “tubulin code” concept, where PTMs convey specific local functionality to the cytoskeleton during neurodevelopment, and second, as a possible general cell biological mechanism of branch-specific axon or synapse dismantling. The “tubulin code” hypothesis suggests that PTMs modify microtubule function, including stability, dynamics, and interactions ^4, 52^. Polyglutamylation is a modification that is especially pronounced in neurons. By changing tubulin’s charge characteristics, glutamylation can affect the recruitment of MAPs to microtubules ^53, 41, 54^. Accordingly, polyglutamylation has been linked to transport deficits and consequent neurodegeneration ^13, 14, 55, 56^. However, physiological roles have been harder to decipher for this PTM. Indeed, the large set of involved enzymes, which catalyze addition and removal of glutamate residues, and the difficulty to probe them with sufficient spatial and temporal precision in the developing nervous system make such investigations challenging. Our translatome-guided and cell-type sepecific genetic approach, now reveal TTLL1-mediated polyglutamylation of tubulin alpha-4A (**Fig. 2** and **5**) as an important step in neuronal pruning and corroborate the high level of enzymatic specificity of the tubulin polyglutamylation system also in neural development ^13^, in full concordance with the “tubulin code” concept. In contrast, the initiator glutamylase TTLL7, which catalyzes the addition of the first glutamate onto tubulin beta isoforms ^57^, does not seem to be involved in remodeling (**Fig. 3**). Notably, tubulin alpha-4A—while not the most abundantly expressed tubulin isoform—is the only tubulin alpha isoform that shows increasing expression across the neuromuscular remodeling period (**Fig. 5a**). This suggests that transcriptional regulation of tubulin genes and perhaps enzymes might be a global factor that is permissive for efficient polyglutamylation-driven remodeling. A further proposition that remains to be tested on the basis of increasing expression (**Fig. 1e**) is that TTLL5, an alternative “seeder” glutamylase for tubulin alpha ^58^, might act upstream of TTLL1 during remodeling.

Our data contain further examples of the complex interplay both on the transcriptional and the post-translational level between various players in this system. For instance, the motor neuron-specific deletion of deglutamylases (CCP1&6^mnKO^) had different outcomes regarding microtubular mass and polyglutamylation levels in developmental remodeling vs. in adult motor axons (**Extended data, Fig. 4b-e**)—two time windows between which spastin mRNA levels dropped (**Extended data, Fig. 4a**). This drop in polyglutamylation could be reversed during remodeling by combined deletion of CCP1 and spastin (**Fig. 7**), suggesting that microtubule stability and apparent PTM levels are the result of the dynamic balance between “eraser” and “reader” activity. In this model, hyper-activated spastin normally removes polyglutamyl-decorated microtubules faster than this population can be replenished—leading to the observed paradoxical relative loss of polyglutamylated microtubules after ablation of the enzymes that normally remove this PTM.

Polyglutamylation can profoundly vary even between adjacent axon branches that only differ in their subsequent fate ^20^. Together with our data, this suggests that glutamate residues can be added onto microtubules not only with temporal but also with high spatial precision. These locally modified microtubules can then, in turn, recruit and activate special sets of MAPs, including tau, kinesin motors, and severing enzymes, such as spastin ^59, 27, 41, 54^. Our data also reveal that changing one PTM (e.g. via genetic manipulation of TTLL1) can percolate within the cytoskeleton to affect other PTMs, such as acetylation ^37^. Thus, the interplay between various PTMs results in a complex pattern of specialized microtubule subsystems within individual axon branches that determines pruning via regulation of microtubule stability and function. Overall, our work implies that polyglutamylation-based encoding can tag neurites for removal—a new model of selective axon removal that can be tested in other species and models.

Open questions that remain, are which events take place upstream and downstream of the spastin-mediated microtubular destabilization. For instance, how does microtubule breakdown result in the characteristic but often divergent neuritic morphologies that accompany pruning? At the neuromuscular junction, axonal breakdown involves the formation of axon bulbs, followed by piecemeal degradation, resembling apoptotic body shedding ^29, 50^. In contrast, mossy fibers in the infrapyramidal bundle and axons in remodeling neurons in the mushroom body seem to rather undergo fragmentation ^50, 60^, while spines are typically absorbed without a morphological trace ^61^. Interestingly, many morphogenic processes in neurites are more commonly associated with actin dynamics rather than with microtubule breakdown ^62, 63^. However, the switch between growth and retraction of neurites is known to involve an interplay of actin and microtubules^64^. Similarly, the balance between the longitudinally rigid co-axial microtubules vs. circumferentially-running actin striations might normally prevent axons from beading^65, 66^. Thus, the idea that polyglutamylation-triggered microtubule loss could trigger shape changes that are seen during pruning is supported by existing evidence, while how the subsequent steps of axonal disintegration followed by glial engulfment are triggered remains to be resolved.

Another key feature of some forms of neuronal remodeling is activity-dependence ^50, 67^. Also here it remains overall unclear how altered activity patterns translate into morphogenic change. In our experiments, chronic α-bungarotoxin blockage of neuromuscular synapses (**Fig. 8**) changed the polyglutamylation levels in presynaptic motor axons. However, it remains elusive, which factor in the enzymatic equilibrium that governs polyglutamylation levels is affected and how the enzymes that mediate or interpret polyglutamylation are regulated by the transcellular signaling that underlies synapse elimination ^64^. For example, spastin phosphorylation might regulate its binding to microtubules or protein stability, although it is not clear how this would impact severing ^68^. For TTLLs and CCPs, information about how their enzymatic activity is regulated is even sparser ^37, 69^. Indeed, thus far no link between regulatory mechanisms that control polyglutamylation and the signaling pathways that are presumed to execute neurite remodeling ^50^ has been made. Our prior work provides a possible feedback mechanism, by which the relevant enzymes themselves might be provided by microtubule-dependent transport, which we have shown to stall early in remodeling ^20^ and influence other signaling events, such as myelination ^49^. Thus, changes in transport could alter the local equilibrium between the addition and the removal of glutamate residues, with spastin acting as a local, branch-specific detector.

Indeed, our work strengthens the long-sought link between developmental neurite remodeling and axon degeneration ^50^. Many of the enzymes of the remodeling mechanism that we describe here are linked to neurodegenerative disease: Spastin to hereditary spastic paraplegia ^8^, CCP1 to infantile-onset neurodegeneration ^70^, and Tuba4a to motor neuron disease ^41^. A unifying mechanism could involve alterations on microtubule dynamics, followed by altered organelle and enzyme transport, which again could feed back to cytoskeletal stability ^71^. Thus, developmental neurite remodeling could be instructive for developing subtler strategies to target microtubules in settings of axon degeneration by focusing drug development on polyglutamylation.

## Acknowledgments

We thank Emily Spaulding and Rob Burgess (The Jackson Laboratory, Bar Harbor, ME, USA) for providing the RiboTag protocol; Julien Gagneur (Technical University Munich, Computational Molecular Medicine, Munich, Germany) for advice on RNA sequencing analysis; Leanne Godinho for critically commenting on the manuscript and Adrian Marti Pastor for advice on software. For excellent technical assistance, we thank Kristina Wulliman, Sabine Brummer and Yvonne Hufnagel; for animal husbandry, we acknowledge Manuela Budak, Nebahat Budak and Matea Korica; we also thank Kristine Kellermann for veterinarian consultancy and Sebastian Berger for IT-support.

## Author contributions

Conceptualization AG, AZ, CJ, TM, MSB; data curation AG, AZ; methodology MB, TM, MSB; funding acquisition TM, MSB; investigation AG, AZ, MW, MR; resources KAZ, MM, TJH, CJ, SE, MK, TM, MSB; supervision CJ, SE, MK, MB, TM, MSB; visualization AG, AZ; writing – original draft AG, TM, MSB; writing – review & editing all authors.

## Experimental Procedures

### Mouse lines, husbandry and genotyping

In all experiments, mice from both sexes were included. Animals were housed in individually ventilated cages with food and water ad libitum. All animal experiments conform to the regulations by the local authorities (e.g., Government of Upper Bavaria). Experimental animals were kept together with littermates.

Motor neuron knockout specific mice were generated by crossbreeding conditional knockout mice to ChAT-IRES-Cre ^32^ mice, expressing Cre-recombinase under the ChAT-promotor (Jackson #6410). ‘Ribotagging’ of motor neurons was conducted in ChAT-IRES-Cre crossbred to homozygous Rpl22^HA^ ^33^ mice (RiboTag^^flox/flox^^; Jackson, #11029) and conditional Spast^^flox/flox^^ (**Fig. 1**, **Fig. 5** and **Extended data, Fig. 1** and **3**). Glutamylases (TTLL1 or TTLL7) or deglutamylases (CCP1 and CCP6) were deleted in motor neurons by crossbreeding conditional mutants of CCP1 ^72^ &6 ^14^, TTLL1 ^14^ or TTLL7 ^13^ mice (gift from Dr. C. Janke, Institut Curie, Orsay) to ChAT-IRES-Cre ^32^ animals. All animals were CCP1&6, TTLL1 or TTLL7 homozygous; experimental animals – CCP1&6^mnKO^, TTLL1^mnKO^ or TTLL7^mnKO^ – were ChAT-IRES-Cre-positive, whereas littermate controls were ChAT-IRES-Cre-negative, named hereafter CCP1&6^mnWT^, TTLL1^mnWT^ or TTLL7^mnWT^. Thy1-YFP ^73^ transgenic mice (cytoplasmic YFP in all motor neurons, Jackson #3709) were used to assess pruning speed in cross-breeding to CCP1&6^mnKO^, TTLL1^mnKO^, TTLL7^mnKO^ compared to their littermates CCP1&6^mnWT^, TTLL1^mnWT^, TTLL7^mnWT^. Microtubule dynamics was visualized and analyzed by Thy1*-*EB3-YFP ^38^ transgenic animals crossed to either CCP1&6^mnKO^, TTLL1^mnKO^ or TTLL7^mnKO^; CCP1&6^mnWT^, TTLL1^mnWT^ or TTLL7^mnWT^ were littermate controls. Conditional knockout of CCP1 ^72^ and Spastin ^20^ in motor neurons, monitored by ROSA-CAG-TdTomato reporter ^74^ (Ai14; Jackson; #7914; CCP1^^flox/flox^^ X Spast^^flox/flox^^ X TdTomato) was generated by injection of viral vectors encoding Cre-recombinase. Block of neurotransmission was conducted and analyzed in constitutive spastin knockout ^20^ (Spast^KO^) mice (homozygous Spast^KO^ vs controls Spast^WT^) and C57BL/6N (Charles River, Strain Code 027) controls injected with α-BTX. Spine density measurements, hippocampal pruning analysis was performed on TTLL1 constitutive knock-out ^14^ (named here TTLL1^KO^; gift from Dr. C. Janke, Institut Curie, Orsay, France) backcrossed to CD-1 (Charles River, Strain Code 022) for three generations. Cytoskeletal analysis and pruning speed were analyzed in homozygous tubulin alpha-4A knock-in ^41^ (Tuba4a^KI/KI^) mice (kindly provided by Dr. M. Kneussel, ZMNH, Hamburg, Germany), and negative, thus wildtype, littermates (Tuba4a^WT/WT^) served as controls. Genotyping of the Tuba4a^KI^ was performed as published ^41^.

Genotyping was performed as previously described ^20, 49^ (for the Tuba4a^KI^ see above). Briefly, genomic DNA was extracted from biopsies using a one-step lysis (lysis buffer in mM: 67 Tris, pH 8.8, 16.6 (NH_4_)_2_SO_4_, 6.5 MgCl_2_, 5 β-mercaptoethanol, 10 % Triton-X-100, and 50 μg/ml Proteinase K; incubation at 55 °C for 5 h, followed by inactivation step 5 min at 95 °C). PCR was performed with GoTaq Green Master Mix (Promega; #M7121) following a standard protocol, and then DNA was separated on a 1.5– 2 % agarose gel. Primer sequences are available upon request.

### Ribosomal pull down and RNA sequencing

Spinal cords were dissected from ChAT-IRES-Cre X Rpl22^HA^ X Spast^flox^ mice (see Mouse lines, husbandry and genotyping above). The samples were kept at −80 °C until pull down of labeled ribosomes described previously ^33^ (see also http://depts.washington.edu/mcklab/RiboTag.html). Spinal cords were homogenized (buffer in RNase free water in mM: 50 TrisCl, pH7.5, 100 KCl (Sigma #P9541), 12 MgCl_2_ (Sigma #63068), 1 % NP-40 (Roche, #11332473001), 1 X Dithiothreitol (DTT, Sigma #646563), 1 X Protease Inhibitors (Sigma #11697498001), 1 mg/ml Heparin (Sigma #H3393), 100 µg/ml cycloheximide (Sigma #C7698), RNAseOUT (Thermofisher #10777019). The mRNA-ribosome complex was precipitated using a polyclonal HA-antibody (Sigma #H6908) and Dynabeads Protein G (Life Technologies #10004D) as previously described. Ribosome-bound mRNA was isolated with RNeasy Plus Micro Kit (Qiagen #74034) per manufacturer instructions (not precipitated fraction was used as “input” spinal cord control). Prior sequencing, RNA quantity and integrity (RIN > 8.5) was controlled with a Bioanalyzer (Agilent RNA 6000 Nano) and rRNA was depleted. Motor neuron mRNA was sequenced in pair-end using an Illumina HiSeq4000 Kit, at a depth of ∼40 million reads per sample (Institute of Neurogenomics, Juliane Winkelmann, Helmholtz Munich, Germany). The raw sequencing data (fastq files) were aligned to the mm9 mouse genome and read counts were extracted with HTSeq-count software with option “intersectionStrict”. Lowly expressed RNAs (< 10 read counts were excluded and differential gene expression analysis was performed with DESeq2 software package ^75^. A likelihood ratio test was performed on the timeline analysis (P5, 7, 9, 11, 14). Graphs were generated using ggplot2 from custom ‘R’ statistical software scripts ^76^.

### Reverse Transcription (RT)-qPCR

All following steps were conducted on ice. cDNA was obtained according to manufacturer’s instruction. Briefly, 100 ng ‘ribotagged’ motor neurons mRNA, 100 µM random hexamer primers (Roche # 11034731001) and 1 μl RNAsin Plus Inhibitor (Promega #N2615) were dissolved with water in 14 μl total volume. The mixture was incubated for 5 minutes at 70 °C, and 10 minutes on ice. 5 μl of M-MLV Reverse Transcriptase Buffer (Promega #M531A), 10 mM NTPs mixture and 200 U of MMLV (Promega #M170A) were added and the mixture incubated for 1.5 h at 37 °C. cDNA was purified with a QIAEX II Gel Extraction Kit (Qiagen #20021) as follows: The obtained cDNA was mixed with 2 μl QiaEX-II suspension (Silica-Matrix) and 80 μl QX-I buffer, incubated for 20 minutes at 25 °C at 1,000 rpm and centrifuged at 13,000 x g for 2 minutes. The supernatant was discarded and the matrix was washed twice with 90 μl PE buffer and centrifuged at 13,000 x g for 2 minutes. The supernatant was removed and the pellet dried for 10 minutes with lid open at 300 rpm shaking at 25 °C. The cDNA was eluted with 20 μl elution buffer (1 mM Tris, pH 8.5), and incubated at 25 °C with 1,000 rpm for homogenization. After centrifugation at 13,000 x g for 2 minutes the supernatant containing the clean RNA was collected.

All qPCR reactions were performed with a LightCyler 1.3 Real-Time PCR system (Roche S/N: 140 6143). The MgCl_2_ concentration (1-3 mM) and the annealing temperature were optimized for each primer pair and confirmed through primer efficiency curves 40. Primer sequence were (5’ to 3’): Chat: TTCTAGCTGTGAGGAGGTGC and CCCAAACCGCTTCACAATGG, GFAP: TCGCACTCAATACGAGGCAG and TTGGCGGCGATAGTCGTTAG. For each reaction, a final concentration of 1 ng/μl cDNA was mixed with 2 μl FastStart DNA Master SYBR Green I (Roche #03 003 230 001), 0.5 μM forward and reverse primer, 1-3 mM MgCl_2_ and water to a total volume of 20 µl. Samples were run at least in triplicates. For each primer pair, controls omitting either template or reverse transcriptase were included.

### Immunofluorescence

The whole thorax was fixed in 4 % PFA for 1 h in 0.1 M phosphate buffer (PB) on ice and the triangularis muscle was dissected and extracted as described previously ^20, 31, 36, 49, 77^. Prior to application of primary antibodies, muscles were incubated in 5 % CHAPS (Carl Roth #75621-03-3) in 0.1 M PB for 1h at 37 °C. Primary antibodies (see details below) were diluted in blocking solution (5 % BSA (Sigma #A7030, 0.5 % Triton-X in 0.1 M PB or 3 % BSA, 0.5 % Triton-X and 12 % NGS (Abcam #ab7481) in 0.1 M PB) and incubated at 4°C overnight for postnatal muscles, 3 days when adult tissue or anti-acetylated tubulin antibody was used. The following primary antibodies were used in this study: anti-tubulin beta-3 conjugated to Alexa Fluor 488 (BioLegend; AB_2562669; mouse IgG2a, 1:200), Alexa Fluor 555 (BD PharMingen; #560339; mouse monoclonal, 1:200), or Alexa Fluor 647 (BioLegend; AB_2563609; mouse IgG2a, 1:200), anti-GT335 (Adipogen, Mouse IgG1, 1: 200), anti-polyE (Adipogen, AG-25B-0030, Rabbit IgG, 1:1,000), anti-acetylated-tubulin (Abcam, 611B1, mouse IgG2b, 1:1,000), anti-neurofilament heavy polypetide (Abcam, ab4680, chicken, 1:500), anti-βmonoE (rabbit, IgG, gift from Carsten Janke, 1:500). Muscles were washed in 0.1 M PB, incubated for 1 h at room temperature with corresponding secondary antibodies coupled to Alexa Fluor 488, Alexa Fluor 594, or Alexa Fluor 647 (Invitrogen; rabbit: #A-11070, #A-11072, #A-21246, #A-32790; mouse: #A-11005; chicken: #A-11042; #A-21449; goat: #A-11058) and washed again in 0.1 M PB. Muscles or sections were mounted in Vectashield (Vector Laboratories) or Prolong-Glass (Thermo Fisher). Image stacks were recorded at a confocal microscope (Olympus; FV1000 or FV3000) equipped with ×20/0.8 NA and ×60/1.42 NA oil-immersion objectives (Olympus).

For immunostaining of spinal cord sections (**Fig. 1**), mice were transcardially perfused with PBS and ice-cold 4 % PFA. Spinal cords were dissected and post-fixed overnight with 4 % PFA. 60 µm thick vibratome sections were then immunostained with antibodies against Choline Acetyl-Transferase (ChAT) (anti-ChAT, Novus Biologicals, goat, NBP1-30052, 1:10) and HA (anti-HA, Sigma-Aldrich, H6908, 1:50), for 2 days over night. Spinal cord sections were washed in 1 x PBS, incubated for 1 h at room temperature with corresponding secondary antibodies coupled to Alexa Fluor 488 and Alexa Fluor 594 (see above). The antibodies for spinal cord staining were diluted in blocking solution prepared with 5 % Normal Donkey Serum (Millipore #S30-M) and 0.5 % Triton-X in 1 x PBS.

Immunostaining of brain sections (**Fig. 4**) was carried out as described previously ^40^. Briefly, mice were transcardially perfused with PBS and ice-cold 4 % PFA in PBS and brains were post-fixed in 4 % PFA in PBS overnight, embedded in paraffin blocks and cut at 7 µm thick coronal sections. Prior staining, antigen retrieval (Citrate-Based Antigen Unmasking Solution, Vector Laboratories) and blocking (1 % BSA, 0.2 % Tween20) was performed. For IPB, visualization sections were incubated overnight at 4 °C with anti-calbindin D28K antibody (Swant #CB-38a, 1:300) diluted in blocking solution. Subsequently, samples were washed three times with PBS with 0.2 % Tween 20 and incubated for 2 hours at room temperature with fluorescent secondary antibody coupled to Alexa Fluor 594 or Alexa Fluor 488 (Life Technologies, #A-11037; A-11008) in blocking solution, washed three times with PBS with 0.2 % Tween 20 and embedded in Mowiol with 1 µg/ml Hoechst. Images were recording using a confocal microscope (Leica SP8 and Leica Stellaris).

### DiOlistics labeling of hippocampal granule cells and dendritic spine density analysis

For visualization of dendritic spines in TTLL1^KO^ brains a DiOlistic approach was used as described previously ^40^. In brief, mice were perfused with 20 ml of 4 % PFA/PBS, and brains were isolated and post-fixed in 4 % PFA/PBS for 30 minutes. Then, brains were washed in PBS for 30 minutes and incubated in 15 % sucrose for 30 minutes, followed by 30 minutes in 30 % sucrose. 250 µm thick coronal slices were cut in vibratome, washed in PBS and incubated for 5 minutes in 15 % sucrose and subsequently in 30 % sucrose. The solution was removed and DiI was introduced into the brain slices by DiI-labeled tungsten particles (prepared as previously described ^40^ with the use of a Gene Gun helium-powered system (Bio-Rad) and 120 Psi pressure. After the labelling, slices were washed in PBS to remove residual tungsten particles and kept for 30 minutes in PBS in the dark to let the dye diffuse. Slices were mounted onto a glass slice in 0.5 % n-propyl gallate/ 90 % glycerol/PBS (NPG) and the next day imaged by a confocal microscope (Leica Stellaris). Analysis of the dendritic spine density was performed in 3 different TTLL1^WT^ and TTLL1^KO^ brains (at least 10 labelled granule cell dendrites were analyzed per brain).

### Virus production

AAV9-hSyn-iCre was produced as outlined previously ^78^. In brief, HEK293-T cells were seeded in 10-tray Cell Factories (Thermo Fisher Scientific) 24 hours before transfection, allowing the cells to reach 80-90 % confluence. Subsequently, 420 μg of the vector plasmid dsAAV-hSyn-iCre and 1.5 mg of the helper plasmid pDP9rs, generously provided by Roger Hajjar (Phospholamban Foundation, Amsterdam, The Netherlands), were introduced into the cells by polyethyleneimine-mediated transfection (Polysciences, 24765 Warrington, PA). Cells were harvested after 72 hours, followed by lysis and benzonase treatment. To purify the AAVs, ultracentrifugation on an iodixanol density gradient (Progen, OptiPrep) was applied. Buffer exchange from iodixanol to Ringer’s lactate buffer was then carried out using Vivaspin 20 columns (Sartorius, VS2042, Göttingen, Germany). To obtain high virus titers, the virus content of two 10-tray Cell Factories was pooled and concentrated. Real-time qPCR with SYBR Green Master Mix (Roche) was performed to evaluate AAV9 titers.

### Neonatal AAV9 or α-BTX injections

Viral vectors were injected into neonatal pups according to previously published protocols ^20, 49, 79^. In short, P3 pups were briefly anesthetized with isofluorane (Abbott) and injected with 3 μl of AAV9-hSyn-iCre (titer 1 x 10^13^ to 2 x 10^14^) into the right lateral ventricle using a nanoliter injector (World Precision Instruments; Micro4 MicroSyringe Pump Controller connected with Nanoliter 2000) attached to a fine glass pipette (Drummond; 3.5”, #3-000-203-G/X) at a rate of 30 nl/s. The injection was guided by ultrasound (Visualsonics, Vevo® 2100). 0.05 % (wt/vol) trypan blue was added to the viral solution for visualizing the filling of the injected ventricles. Whole litters were injected, and pups were allowed to recover on a heating mat before the litter was returned to their mother into the home cage and sacrificed on P9 for experiments.

For analysis, Cre-mediated deletion in conditional CCP1 and Spast knockout (CCP1^^flox/flox^ 72^ X Spast^^flox/flox^ 20^) was verified by a robust expression of the TdTomato reporter allele ^74^ (homozygous) in motor neurons of the triangularis sterni muscle. According to TdTomato fluorescence intensity in axons, recombination was assumed, when fluorescence was in the upper quantile and negative when they belonged to the lowest quantile. All other axons were excluded from analysis.

To block neurotransmission in the triangularis sterni muscle of C57BL/6N (Charles River, Strain Code 027) or Spast^KO 20^ mice, 1 µl of 50 µg/µl α-BTX conjugated to Alexa Fluor 594 (Invitrogen; B13423) was injected with a needle unilaterally in the thorax on P7, as previously described ^49^. In some cases, a post-hoc stain of the triangularis sterni muscle with anti-Neurofilament (see above) or α-BTX Alexa Fluor 488 or 594 further verified the degree of labeling with injected α-BTX conjugated Alexa Fluor 594 and absence of denervation (> 100 neuromuscular junctions (NMJs) per mouse, n = 3 mice). Immunostaining and quantification were conducted as described above with anti-tubulin beta-3 and anti-polyE antibodies.

### Live imaging of EB3 comet densities in nerve-muscle explants

EB3 comet imaging was performed on acute nerve-muscle explants as previously described ^20, 38^. In brief, the thorax of euthanized mice was obtained by removing the skin over the rib cage, severing the ribs close to the spinal column and dissecting the diaphragm. Further dissections were done under oxygenated, ice-cold Ringer’s solution (in mM: 125 NaCl, 2.5 KCl, 1.25 NaH_2_PO4, 26 NaHCO_3_, 2 CaCl_2_, 1 MgCl_2_, and 20 glucose, oxygenated with 95 % O_2_/ 5 % CO_2_) to remove thymus, pleura, lung and pectoral muscles over the rib cage. The explant was fixed with insect pins (Fine Science Tools; 26001– 25, 0.25 mm) on a Sylgard-coated 3.5 cm petri dish, with the inside of the thorax facing the objective. Imaging was performed under continuous and steady perfusion with warmed oxygenated Ringer’s solution and kept at physiological temperatures 33-36 °C by a heated stage connected to an automatic temperature controller (Warner Instruments; TC-344C). Total imaging time on explants did not exceed 2 hours. Live imaging was carried out with an epifluorescence microscope (Olympus BX51WI) equipped with ×20/0.5 NA and ×100/1.0 NA water-immersion objectives, an automated filter wheel (Sutter Instruments; Lambda 10–3), a charge-coupled device camera (Visitron Systems; CoolSnap HQ2), controlled by μManager version 1.4 ^80^. A total of 200 frames were acquired per movie, at a frequency of 0.5 Hz and an exposure time of 500 ms using a YFP filter set (F36-528; AHF Analysentechnik).

### Data analysis

ImageJ/FiJi ^81^ (http://fiji.sc) was used to determine pruning speed, by counting the number of innervating terminal branches ending on each α-BTX-stained neuromuscular junction. For quantification of immunostainings three background subtracted areas (ROIs; measured for mean gray value) were averaged per single axon. Each ROI was obtained from a single optical section, as previously described ^20, 49^. IPB length was quantified using the ratio of IPB length to the length of the CA3 as described previously ^82^. To quantify EB3 comet parameters, out-of-focus frames were deleted manually from movies and aligned with the TurboReg plugin with the parameter Rigid-body and “Quality” set on “Accurate” and comet trajectories were manually analyzed using the MTrackJ-plugin (developed by E. Meijering, Biomedical Imaging Group, Erasmus Medical Center, Rotterdam). Only EB3 comets that appeared for at least three consecutive frames were considered. Neurolucida software (MBF Bioscience) was used to perform dendritic spine segmentation.

For image representation, maximum intensity projections were generated from confocal image stacks with ImageJ/Fiji ^81^ and then further processed in Adobe Photoshop. All analyses were performed with the experimenter blinded to the treatment or genotypes during imaging and scoring.

### Statistics

Statistical analysis and visualization were performed using ‘R’ ^76^ or OriginLab software (origin.lab). All experiments included at least 3 biological replicates. Normality was tested and outliers were removed using the Grubbs test. A two-tailed Students t-test was performed if data passed the normality test, otherwise Mann-Whitney test was used for non-parametric data. P < 0.05, indicated as *, was considered significant. P < 0.01 is **, P < 0.001 is ***, and P < 0.0001 is ****. Bars – on the left – show mean + SEM. Violin plots – on the right – depict data distribution, scaled based on constant maximum width (graphs are cut at 90 % level of the dominant violin).

