## Supplemental figures for "Polyglutamylation of microtubules drives neuronal remodeling"

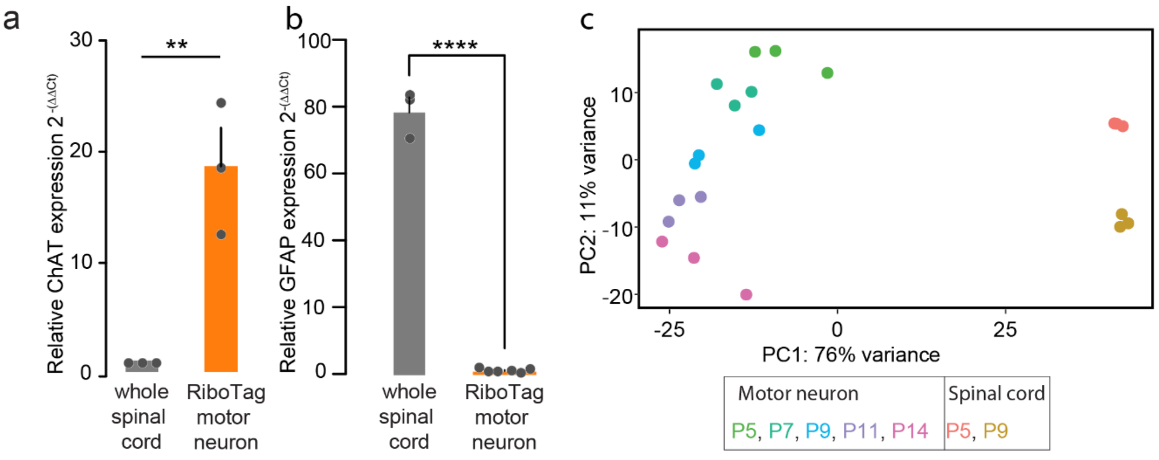


d

GO "Biological Process" in motor neuron vs spinal cord at P9

| GO term | Description | Bonferroni |
|---|---|---|
| 0000902 | cell morphogenesis | 6.25e-11 |
| 0006812 | cation transport | 2.03e-10 |
| 0035295 | tube development | 1.73e-9 |
| 0030198 | extracellular matrix organization | 3.97e-9 |
| 0048666 | neuron development | 4.03e-9 |
| 0031175 | neuron projection development | 4.13e-9 |
| 0061564 | axon development | 1.97e-8 |

e

GO "Cellular Component" in motor neuron vs spinal cord at P9

| GO term | Description | Bonferroni |
|---|---|---|
| 0045202 | synapse | 9.21e-6 |
| 0043209 | myelin sheath | 1.73e-5 |
| 0030424 | axon | 3.39e-4 |
| 0098793 | presynapse | 7.26e-4 |
| 0044297 | cell body | 2.12e-3 |
| 0015630 | microtubule cytoskeleton | 2.64e-3 |
| 0031252 | distal axon | 1.63e-2 |

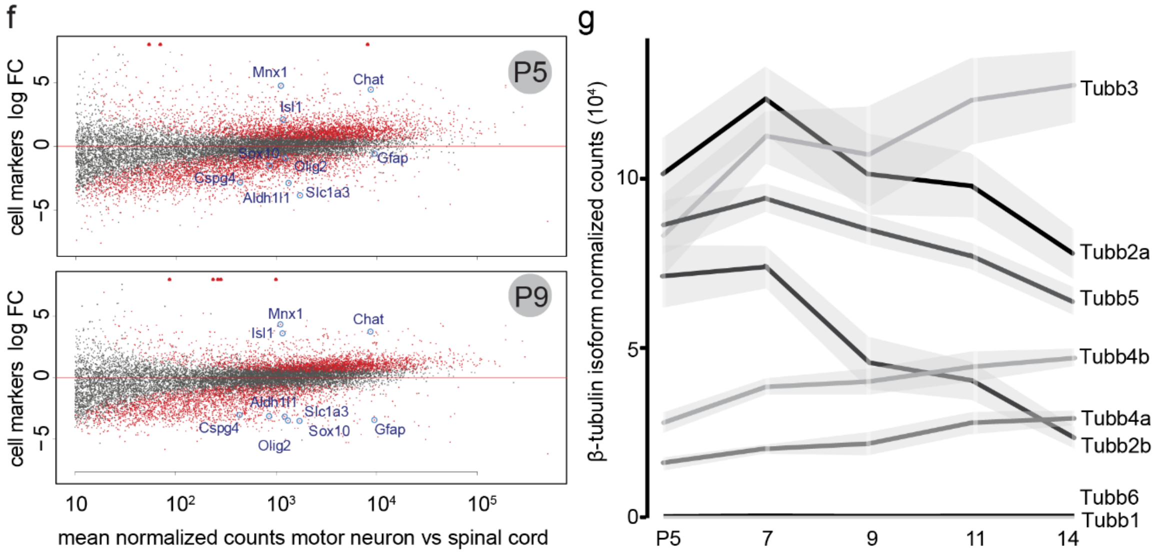


**Extended data; Figure 1. Confirmation of motor neuron specificity of the RiboTag approach**

**(a)-(b)** RT-qPCR on whole spinal cord and motor neuron RiboTag (Rpl22$^{HA}$ + IP) samples, of (a) Choline Acetyl Transferase (ChAT) and (b) Glial Fibrillary Acidic Protein (GFAP). Results were calculated according to 2-Δ Δ Ct method, using primers of a tested efficiency ≥ 90%. The amplification threshold cycle (Ct), was the average of 2 or 3 technical replicates from each biological sample (n ≥ 3 animals per group). **(c)** PCA plot of mRNA samples from spinal cord (sc) and mRNA from IP from motor neurons (mn). Individual dots represent a single animal, color coded by age. **(d)** GO category "Biological Processes" generated using top 500 most differentially expressed genes by comparing motor neuron against spinal cord at P9 using a cutoff of $p_{adj} \leq 1e\text{-}7$ and $|Log_2\ FC| > 2$. **(e)** GO category "Cellular Component", generated by using top 150 most differentially expressed genes by comparing motor neuron against spinal cord at P9 using a cutoff of $p_{adj} \leq 1e\text{-}7$ and $|Log_2\ FC| > 2$. **(f)** MA plot at P5 and P9 highlights enrichment of motor neuron specific cell markers in $Rpl22^{HA}$+ motor neuron translatome samples, compared to total spinal cord. Genes above the cutoff threshold of FDR ≤ 0.05 and a $|Log_2\ FC| \geq 1.5$ are depicted in red. Genes above the red horizontal line are enriched in pull-down group, genes below the red horizontal line are reduced. **(g)** Graph depicts normalized mRNA counts ($*10^4$) of tubulin beta isoforms across the motor axon remodeling phase (postnatal day (P) 5, 7, 9, 11, 14). Graphs: mean + SEM and data representing single animals as single dots (a,b) or mean ± SEM (g). Mann-Whitney determined significance: **, P< 0.01; ****, P < 0.0001.

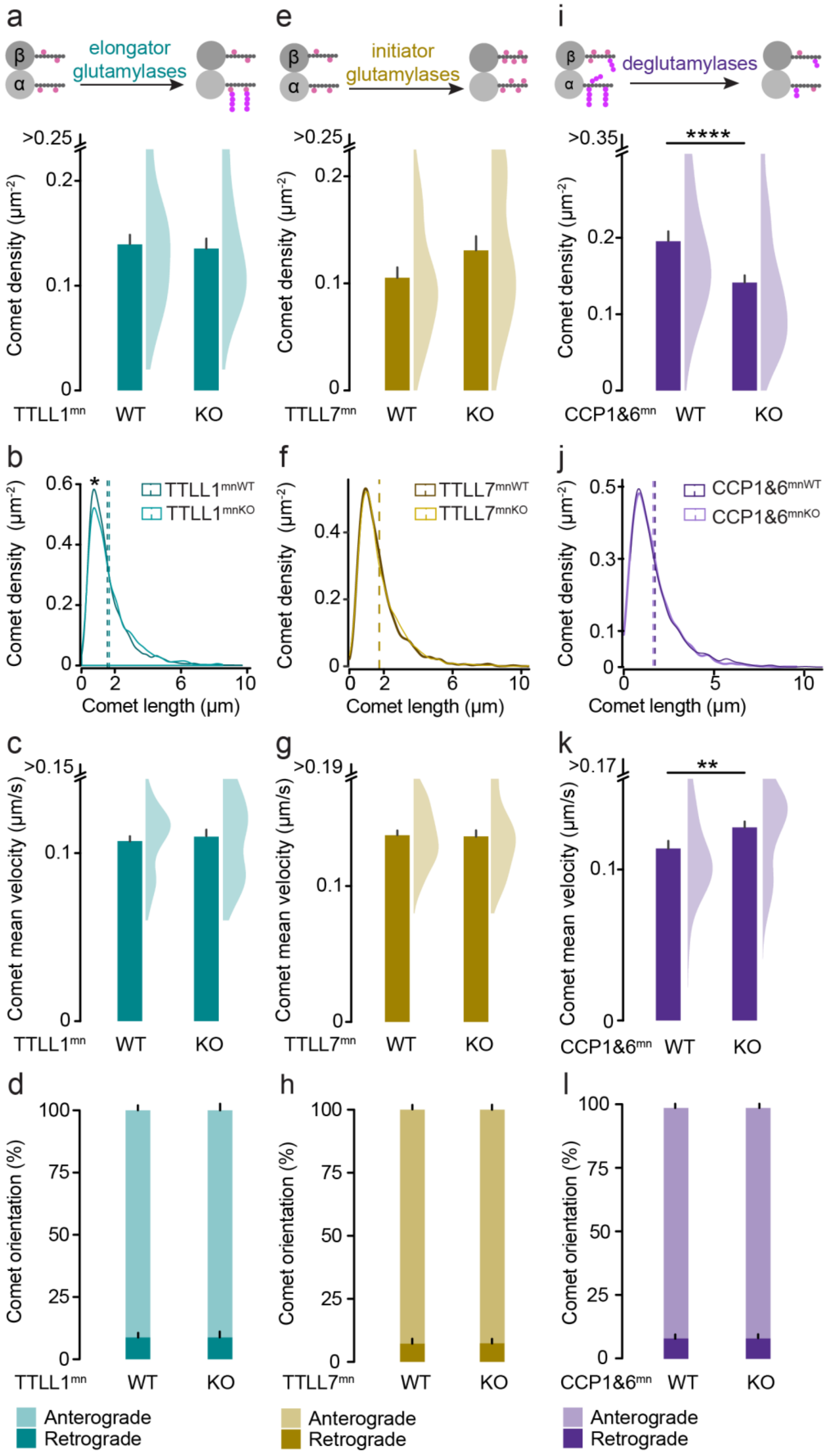

a
elongator glutamylases
β
α
e
initiator glutamylases
i
deglutamylases
>0.25
Comet density (µm⁻²)
TTLL1mn WT KO
>0.25
TTLL7mn WT KO
>0.35
****
CCP1&6mn WT KO
b
TTLL1mnWT
TTLL1mnKO
Comet length (µm)
f
TTLL7mnWT
TTLL7mnKO
j
CCP1&6mnWT
CCP1&6mnKO
c
>0.15
Comet mean velocity (µm/s)
g
>0.19
k
>0.17
**
d
Comet orientation (%)
h
l
Anterograde
Retrograde

**Extended data; Figure 2. Microtubule dynamics in terminal motor axons in explants of TTLL1$^{mnKO}$, TTLL7$^{mnKO}$, CCP1&6$^{mnKO}$ crossbred to Thy1-EB3-YFP**

**(a)-(l)** Based on live imaging of EB3-YFP in nerve-muscle explants of P8-11 TTLL1$^{mnKO}$ (a-d), TTLL7$^{mnKO}$ (e-h) and CCP1&6$^{mnKO}$ (i-l) animals, we analyzed comet density (a,e,i), comet length distribution (b,f,j), mean velocity (c,g,k) and orientation (d,h,l). (a-c: n ≥ 4 animals per genotype, n ≥ 38 axons per genotype; d-f: n ≥ 4 animals per genotype, n ≥ 26 axons per group; g-i: n ≥ 4 animals per genotype, n ≥ 43 axons per group)

Graphs: mean + SEM (left) and data representing axons as half violin (right) (a,c,e,g,I,k), mean (dashed vertical line) + data (line) (b,f,j) and mean + SEM (d,h,l). Mann-Whitney test determined significance: *, $P < 0.05$; **, $P< 0.01$; ****, $P < 0.0001$.

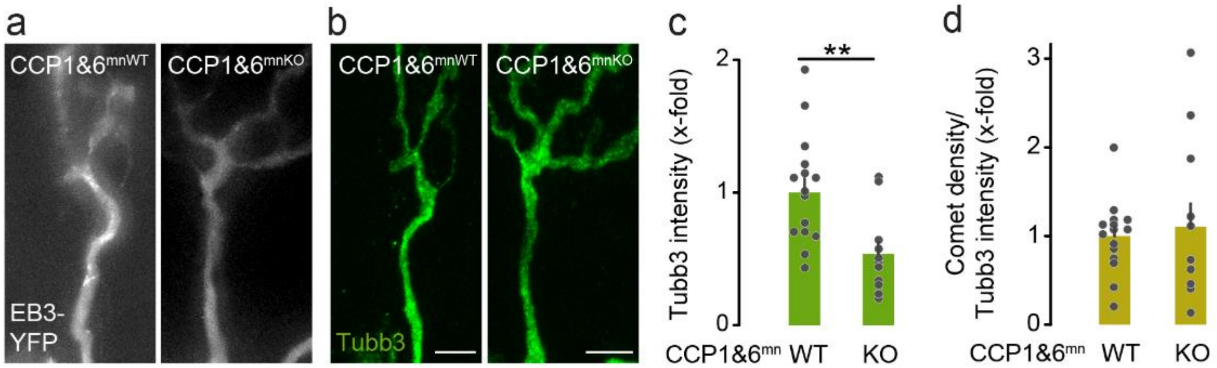


**Extended data; Figure 3. Correlative analysis of EB3-YFP comet to tubulin beta-3 intensity in CCP1&6$^{mnKO}$ crossbred to Thy1-EB3-YFP**

**(a)-(c)** Correlation of EB3-YFP-imaging in explants of P9-11 CCP1&6$^{mnKO}$ and CCP1&6$^{mnWT}$ littermates to post-hoc immunostaining for tubulin beta-3 in the same terminal motor axons.

**(a)** Maximum intensity projection stacks of neuromuscular synapse captured from time-lapse recording (20s) of EB3-YFP comets (gray).

**(b)** Confocal image of the same terminal neuromuscular synapse (represented in j), immunostained for tubulin beta-3 (Tubb3, green).

**(c)** Quantification of Tubb3 intensity and **(d)** quantification of EB3 comets density to Tubb3 intensity ratio.

Graphs: mean + SEM and data representing animals as single dots (c-d). Mann-Whitney test determined significance: **, P< 0.01. Scale bars, 5 µm.

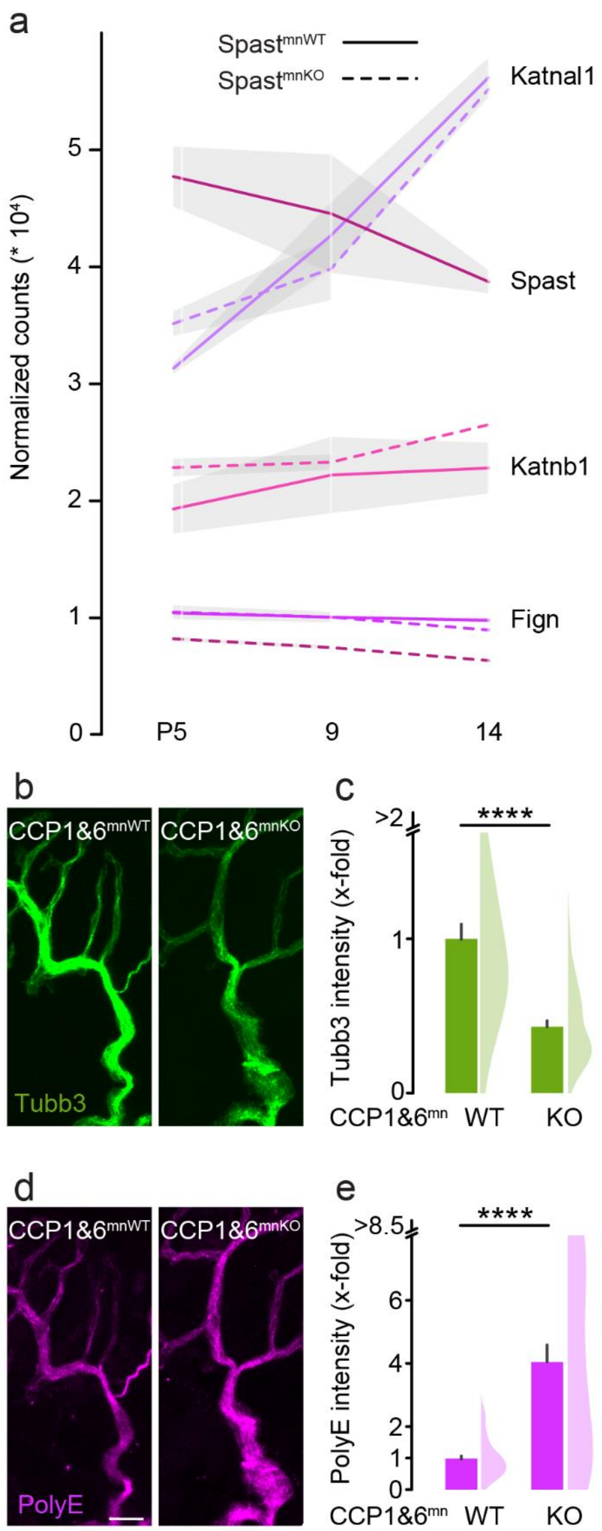


**Extended data; Figure 4. Severing enzymes in motor neuron translatome across neuronal remodeling and microtubule alterations in CCP1&6$^{mnKO}$ adult motor neurons**

**(a)** Normalized read counts (*10$^4$) from translatome analysis of severing enzymes in motor neurons of Spast$^{mnWT}$ or Spast$^{mnKO}$ mice at P5, 9 and 14 (dashed line is spast$^{mnKO}$, regular line spast$^{mnWT}$).

**(b)–(e)** Quantitative immunostainings for tubulin beta-3, neurofilament heavy polypeptide and polyglutamate on triangularis sterni muscles at 6 weeks from CCP1&6$^{mnKO}$ and CCP1&6$^{mnKO}$ control littermates. **(b)** Confocal stack of a neuromuscular synapse depicting tubulin beta-3 staining (Tubb3, green). **(c)** Quantification of Tubb3 intensity in terminal motor axons, normalized to neurofilament heavy polypetide (n = 3 mice per genotype, n ≥ 33 axons per genotype). **(d)** Confocal stack of a neuromuscular synapse depicting polyglutamate chain staining (PolyE, magenta). **(e)** Quantification of polyE intensity in terminal motor axons, normalized to tubulin beta-3 (n = 3 mice per genotype, n ≥ 33 axons per genotype).

Graphs: mean ± SEM (a) and mean + SEM (left) and data representing axons as half violin (right) (b,d). Mann-Whitney test determined significance: ****, P < 0.0001. Scale bars, 5 µm.

### Supplemental video legends

**Extended data; Movie 1. Microtubule dynamism in P9 Thy1-EB3-YFP x CCP1&6$^{mnKO}$ motor neurons**

Time-lapse movie, depicting tracking of EB3 comets (colored lines) visualized in nerve-muscle explants of a crossbreeding of Thy1-EB3-YFP and CCP1&6$^{mnKO}$ (bottom) vs. CCP1&6$^{mnWT}$ (top) at P9.

### Supplemental table legends

**Extended data; Table 1. Gene differential expression between whole spinal cord and RiboTag motor neuron samples**

List of differentially expressed genes at P9 – whole spinal cord vs. RiboTag – translatome, summarizing the Log2 Fold Change (Log$_2$|FC|) of expression between motor neuron and spinal cord.

**Extended data; Table 2. Motor neuron translatome during neuronal remodeling**

Complete list of motor neuron translatome derived genes, summarizing normalized transcript counts from 3 biological replicates including whole spinal cord and RiboTag samples at P5, 7, 9, 11, 14.

**Extended data; Table 3. Gene differential expression between Spast$^{mnWT}$ and Spast$^{mnKO}$**

List of all motor neuron translatome derived genes as well as differentially expressed genes between Spast$^{mnWT}$ and Spast$^{mnKO}$ at P5.
